# Design Principles of Lambda’s Lysis*/*Lysogeny Decision vis-a-vis Multiplicity of Infection

**DOI:** 10.1101/146308

**Authors:** Dinkar Wadhwa

## Abstract

Bacteriophage lambda makes a decision between lysis and lysogeny based on the number of coinfecting phages, namely the multiplicity of infection (MoI): lysis at low MoIs; lysogeny at high MoIs. Here, by evaluating various rationally designed models on their ability a) to make the lytic decision at MoI of 1 and the lysogeny decision at MoI of 2, b) to exhibit bistability at both MoIs, and c) to perform accurately in the presence of noise, it is demonstrated that lambda’s lysis/lysogeny decision is based on three features, namely a) mutual repression, b) cooperative positive autoregulation of CI, and c) cooperative binding of the activator protein, not basal expression, triggering positive autoregulatory loop of CI. Cro and CI are sufficient to acquire the first two features. CII is required to acquire the third feature. The quasi-minimal two-protein model for the switch is justified by showing its qualitative equivalence, except for Cro repression of *pRM*, to the lambda’s gene regulatory network responsible for the decision. A three-protein simplified version of the lambda’s switch is shown to possess all the three design features. Bistability at MoI of 1 is responsible for lysogen stability, whereas bistability at MoI of 2 imparts stability to lytic development post-infection and especially during prophage induction.

## 1. Introduction

While virulent bacteriophages reproduce only through lysis, temperate bacteriophages have an additional perpetuation strategy, namely lysogeny. In lysis, the phage injects its genetic material into the bacterial host, viral genes are transcribed, mRNAs thus produced are translated, and the phage’s genetic material is replicated. Finally, the bacterial cell is lysed, thereby releasing the progeny particles. In lysogeny, the lytic pathway is repressed, the viral genome is integrated into, and replicated with, the bacterial chromosome, and thus the virus exists in its latent state known as prophage. As the teleological explanation goes, the lytic strategy leads to fast multiplication, but it is risky because the viral progenies have to find new hosts which do not already contain lysogenized phages. On the other hand, a lysogenized phage replicates along with its host and hence reproduces slowly as compared to the reproduction through the lysis strategy, but this way the phage safeguards its survival. Should a phage infect a bacterium containing lysogenized phage, lambda repressors (CI) present in the cytosol will suppress the former’ lytic program.

Being one of the simplest organisms to make a genetic decision; that is, between lysis and lysogeny, bacteriophage lambda has been a paradigm for studying gene regulatory networks and developmental decision making between alternative cell fates. Extensive study of the phage’s core genetic network involved in the lysis/lysogeny decision has yielded a complex picture of interconnected positive and negative feedback loops [1]. Two studies, one classic [2] and another molecular [3], have shown that the lysis/lysogeny decision depends on multiplicity of infection (MoI). Avlund et al. [4] analysed Kourilsky’s data [2, 5] and determined the probability of lysogeny at MoI of 1 to be almost zero, at MoI of 2 to be around 0.6960, and at all the higher MoIs to be around 0.9886. Abstractly, this ability of the phage to choose between lysis and lysogeny based on the number of coinfecting phages is but a form of quorum sensing occurring inside a bacterium.

One of the first theoretical studies of lambda’s lysis/lysogeny decision with respect to multiplicity of infection considered a model which possessed two proteins, namely Cro and CI, produced from their respective promoters, namely *pR* and *pRM*, and a simplified model of the phage’s gene regulatory network (GRN) which possessed three critical proteins, namely Cro, CI, and CII. Cro and CI had the same promoters as in the two-protein model. CII is under the control of the *pR* promoter and activates the third promoter *pRE*, whose reading frame contains *CI* [6]. In both models, all proteins bind as dimer. For the two-protein model, it was shown that if basal expression of the *cro* gene exceeds that of the *cI* gene, only one stable state exists at low MoI, where Cro ¿¿ CI. Similarly, only one stable state exists at high MoI, where CI ¿¿ Cro. On the other hand, two stable states exist for the intermediate MoIs.

Additionally, it was demonstrated that Lys positive autoregulation and protein dimerization are necessary for producing the switching behaviour. We use the word “switch” or the phrase “switching behaviour” throughout this paper to mean the generation of Lyt-dominant state at low MoI and Lys-dominant state at high MoI. For the three-protein model, the work showed that the transient level of CII increases with MoI, and past a certain threshold, the nonlinearity of CI positive autoregulation gives rise to the switching behaviour in terms of equilibrium values of the three proteins. Although the study demonstrated that a two-protein and three-protein reduced models of lambda’s lysis/lysogeny switch are able to produce the switching behaviour deterministically, it did not determine how the switching behaviour is rooted in the features of the models. For example, the study did not address the question as to what advantage does the additional protein CII confer to the two-protein model, which also is able to produce the switching behaviour.

Avlund et al. constructed all possible motifs comprised of two proteins, namely Cro-like protein (Lyt) and CI-like protein (Lys) [7]. The networks were selected on their ability to stably commit to lysis and lysogeny at MoI of 1 and MoI of 2, respectively, by solving their corresponding dynamical equations deterministically (the “deterministic counting task”). 96% of the networks successful in the deterministic counting task possessed mutual repression motif. In the subsequent work, Avlund et al. tested the two-protein networks successful in the deterministic counting task for their ability to carry out the same task in the presence of noise (the “stochastic counting task”) [8]. It was found out that almost all of the two-protein networks did not pass the stochastic counting task. However, additional CII-like protein significantly improved robustness to noise.

The mean lifetime of CII in the networks passing the deterministic counting task was 0.8 phage replication time (τ*_rep_*). The stochastic counting task selects for CII with shorter half-life: the mean lifetime of CII in the networks carrying out the stochastic counting task reduced to 0.4 τ*_rep_*. The mean lifetime of Lys in the two-protein networks successful in the deterministic counting task was 1.2 τ*_rep_*, and this increased to 4.6 τ*_rep_* in the presence of CII. Thus, the authors argued that Lys, because of having longer mean half-life, reaches its steady state slowly and, as a result, interferes less with CII, which has much shorter half-life, during the commitment phase. Hence, it was argued that the reason for the improved robustness was CII relieving Lys of its role of executing lysogenic commitment, thereby allowing Lys to be independently optimized for lysogenic maintenance.

In this study, conceptualizing the roles of Cro-like protein from first principles, a two-protein quasi-minimal model for the lambda’s GRN is obtained; and additionally, other two-protein models based on certain departures from the quasi-minimal model are constructed. Finally, a three-protein simplified model of the lambda’s switch was constructed too. Together with the constraints of producing a sound quality switching behaviour and possessing bistability at both MoIs, relating the accuracy of the switching behaviour in the presence of noise with its respective model features reveal design principles of the lysis/lysogeny switch.

## 2. Result and Discussion

### 2.1. Quasi-minimal two-protein lysis***/***lysogeny switch

The quasi-minimal two-protein model consists of a Cro-like protein (Lyt) and a CI-like protein (Lys) whose corresponding genes are *lyt* and *lys*. Promoter of *lyt* is constitutive and that of *lys* is positively regulated as they are in the lambda’s GRN. The roles of Lys in the quasi-minimal two-protein model; that is, binding cooperatively to the intergenic region, activating transcription of its own gene and inhibiting transcription of *lyt* gene, are identical to the functions of CI in the lambda’s GRN; that is, binding cooperatively to the intergenic region, activating the *pRM* promoter and repressing the *pR* promoter. Additionally, Lys activates an imaginary downstream pathway which causes lysogenic development.

The role of Lyt was conceptualized from first principles in the following way. At MoI of 1, constitutive expression of *lyt* results in Lyt production to a high equilibrium level. On the other hand, the level of Lys remains low because *pRM* is a very weak promoter [9, 10]. Although the analysis of Avlund et al. [4] showed that lysogeny occurs in around 70% of the cases at MoI of 2, in the present study, for the sake of simplicity, lysogeny is expected to always occur at MoI of 2. Hence, at MoI of 2 the reverse situation should occur: the equilibrium level of Lyt (Lyt_2_) should be much lower than that at MoI of 1 (Lyt_1_), whereas the equilibrium level of Lys (Lys_2_) should be much higher than that at MoI of 1 (Lys_1_). However, that cannot happen because *lys*, being regulated by a very weak promoter, cannot produce Lys to a level sufficient to trigger the autoregulatory loop of Lys. Because the only protein present at MoI 2 to actuate any process is Lyt, we argue from first principles that Lyt should activate promoter of *lys* and suppress that of its own.

Lyt is produced constitutively at MoI of 1 but its level is not high enough to trigger positive autoregulatory loop of Lys, causing the level of Lys to remain low. On the other hand, Lyt is initially produced at a higher rate at MoI of 2, triggering the positive feedback loop, which results in Lys production. Lys, thus present at a high concentration, represses *lyt*, causing Lyt level to come down to a low value. Thus Lyt carries out three functions: 1) activates transcription of *lys*, whose product engenders lysogeny and suppresses the lytic pathway, 2) represses transcription of its own gene, thereby suppressing the lytic pathway (though, as argued below, this interaction is dispensable when kinetic details are ignored), and 3) activates an imaginary downstream pathway which causes lytic development. As explained below, these seemingly paradoxical roles of Lyt; that is, causing lysogeny as well as lysis, is because of it being a proxy for CII, which causes lysogeny, and anti-termination factor Q, which enables transcription of the lytic genes.

Because the expression for the quality of the switching behaviour (the “switch quotient”) takes equilibrium values into account, the degradation rate of Lyt (*X*) can be subsumed into *k*_1_, and the degradation rate of Lys (*Y*) can be subsumed into *k*_3_ and, if present, *k*_4_. Hence, *k*_2_ and *k*_5_ are taken to be unity for all of the two-protein models. This model would henceforth be referred to as 1A_Lyt_Lys.

#### 1A_Lyt_Lys

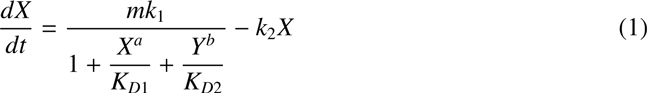

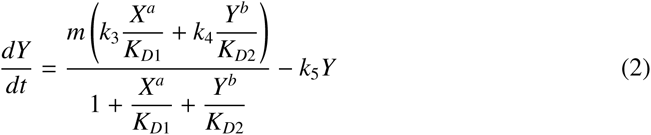

where, *m* is multiplicity of infection; *k*_1_ is the basal expression rate of *lyt*; *k*_3_ and *k*_4_ are the rate constants for transcriptional activation of *lys* by Lyt and Lys, respectively. *K_D_*_1_ and *K_D_*_2_ are the effective equilibrium dissociation constants of Lyt and Lys, respectively (see Methods). In those models in which *lys* has a basal expression, *k*_3_ represents the basal expression rate. The exponents a and b are the Hill coefficients for Lyt and Lys binding, respectively.

### 2.2. Equivalence of the quasi-minimal two-protein model (1A_Lyt_Lys) with the lambda’s GRN

Upon the infection, RNA polymerase transcribes from the early promoters *pL* and *pR* until it encounters the transcription terminators *tL1* and *tR1*, respectively [11]. The *tL1* terminated transcript possesses the *N* gene. The product of the *N* gene is an anti-termination factor that modifies subsequent RNAPs initiating at *pL* and *pR*, so that they are able to transcribe genes that are beyond their terminators, such as *cIII* and *cII* genes, respectively [12, 13]. Such a modified RNAP from promoter *pR* is also able to transcribe through another terminator, namely *tR2*, present upstream of the gene *Q* (see figure 2). Up to this point, the lytic and lysogenic pathways are identical. Commitment to lysis occurs in the following way: extended transcription from *pR* results in transcription of gene *Q*. Q, being an anti-termination factor, causes transcription from the strong constitutive promoter *pR*’ to not terminate, as it would otherwise normally do, at *tR*’, which is present at about 200 bases away from *pR*’. This results in transcription of the rest of the lytic genes downstream of *Q* [14], causing the cell to commit to lysis. CIII protein has an indirect role in causing lysogeny, that is, by preventing degradation of CII by the bacterial protease HflB [15, 16]. The (indirect) role of cIII will not be taken into consideration in the present work, as it investigates the decision-making process of lambda through minimalist models.

**Figure 1:**
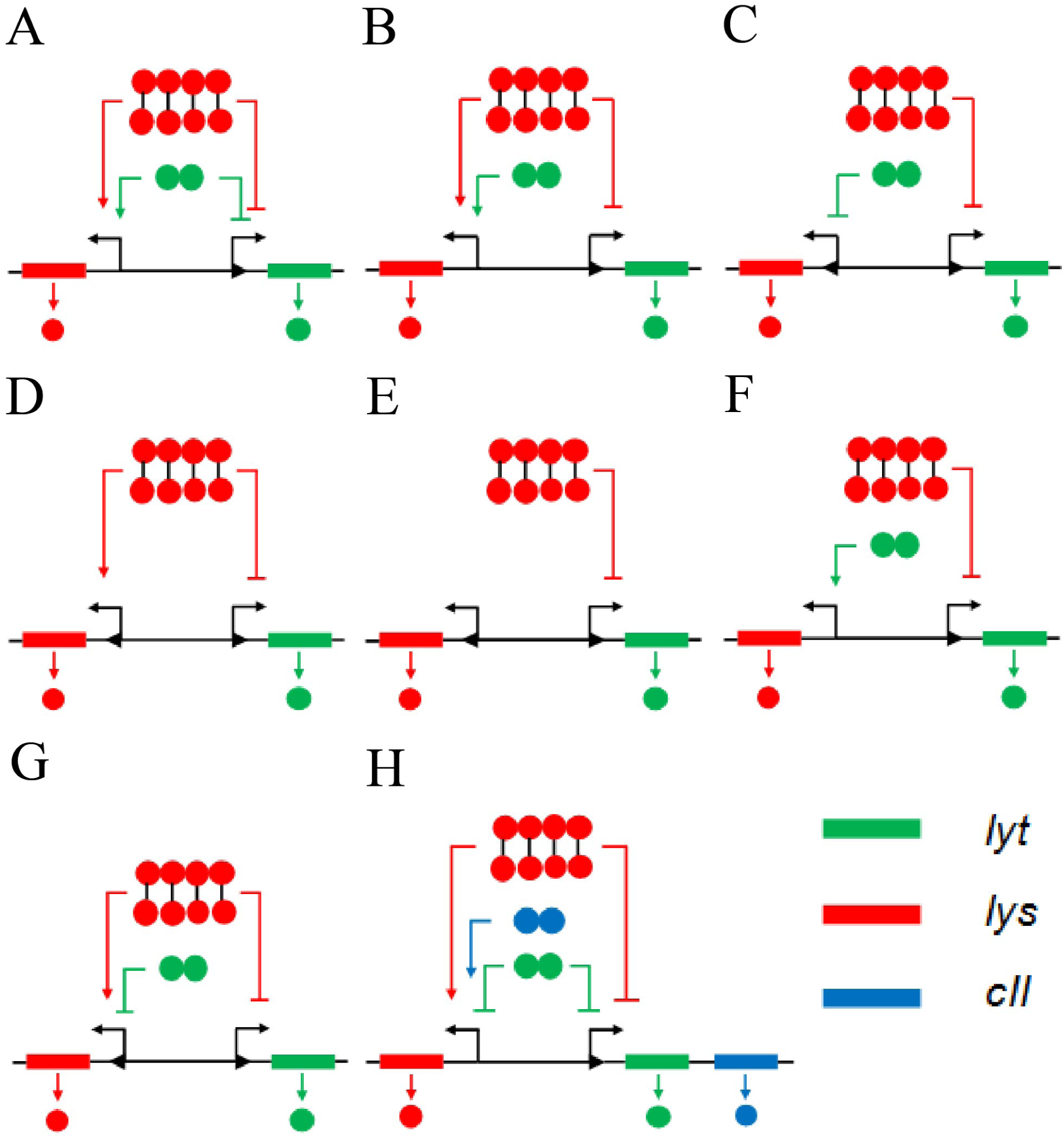
Various two-protein models, and the three-protein model. Lower arrowhead represents basal expression. (**A**) The quasi-minimal model or 1A_Lyt_Lys. (**B**) 1A_Lyt_Lys with self-repression of *lyt* removed, or 1B_Lyt_Lys. (**C**) Mutual repression or 2_Lyt_Lys. (**D**) 3_Lyt_Lys. (**E**) 4_Lyt_Lys. (**F**) 5_Lyt_Lys. (**G**) 6_Lyt_Lys. (**H**) A three-protein simplified version of the lambda’s switch or Lyt_Lys_CII.

**Figure 2:**
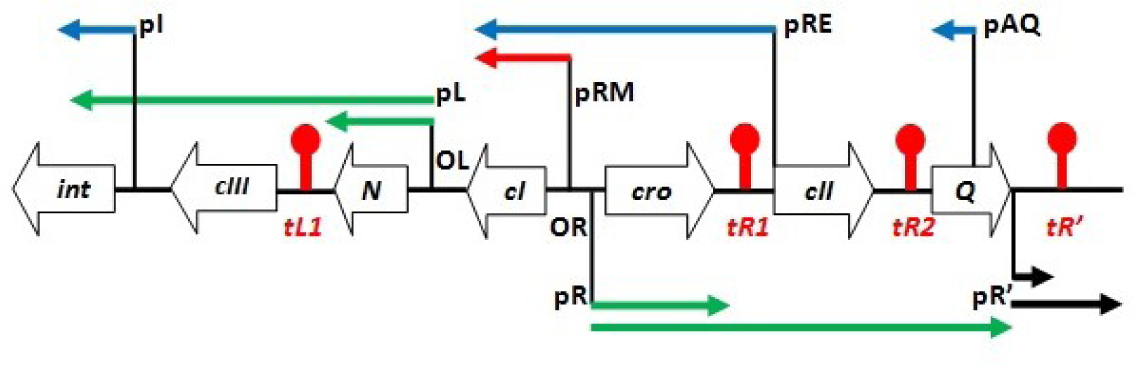
Gene and transcription map of lambda. Transcripts produced from the early promoters, namely *pL* and *pR*, are depicted as green arrows. The black arrow represents transcripts generated from the late promoter *pR*’. Transcripts produced from CII-activated promoters, namely *pI*, *pRE*, and *pAQ*, are shown as blue arrows. Transcripts from *pRM* are shown as red arrow. The transcription terminators, namely *tL1*, *tR1*, and *tR2*, are depicted by a red club.

Commitment to lysogeny occurs when CII is produced to a concentration which is sufficient to activate transcription from three promoters, namely *pI*, *pRE*, and *pAQ*. The *pI* promoter controls transcription of *int* gene, whose product, the Int protein, is required for the integration of the phage genome into the bacterial chromosome [17]. Because mRNAs produced from *pRE* contain the gene *cI*, activation of this promoter results in CI production [18]. CII inhibits commitment to the lytic pathway by activating transcription from *pAQ*, which is located within the *Q* gene in the opposite polarity [19, 20]. Being antisense to (a part of) *Q* mRNA, transcripts produced from *pAQ* hybridize with *Q* mRNAs. This prevents translation of *Q* mRNAs, whose translated product is essential for the commitment to lysis [3].

Because in 1A_Lyt_Lys *lyt* is transcribed from a *pR*-like promoter, Lyt should be functionally equivalent to *pR*-regulated proteins, namely Cro, CII, and Q. Lyt negative autoregulation is equivalent to Cro repression of *pR*. Because *cII* and *Q* are both regulated by *pR*, Lyt self-repression is also equivalent to CII inhibition of Q’s production, that is, by activating transcription from *pAQ*. Lyt activation of *lys* expression is equivalent to CII activation of *pRE*, which results in the synthesis of CI. CII activation of *pI*, whose product is required for lysogeny, is modeled by Lyt activating promoter of *lys*, whose product activates the imaginary downstream pathway that causes lysogeny. Thus, promoter of *lys* is equivalent to those promoters whose activation causes lysogeny, namely *pRE* and *pI*, and Lys is equivalent to those proteins whose genes are regulated by these promoters, namely CI and Int. If CII is not produced in sufficient amount, *Q* mRNA is translated and the anti-terminator protein Q thus produced causes lysis. The quasi-minimal model incorporates this process in this way: if Lyt concentration does not become high enough to be able to trigger positive autoregulatory loop of Lys, which results in Lys production and consequent inhibition of *lyt*, Lyt is produced to a concentration which is sufficient to cause the commitment to the lytic pathway.

Notably, while the Cro protein in the lambda’s GRN inhibits transcription from the *pRM* promoter, the Cro-like protein (Lyt) activates expression of *lys* in 1A_Lyt_Lys. It will be shown later that inhibition of *lys* (or *pRM*) by Lyt (or Cro) is required for the stability of lytic development post-infection and especially during prophage induction. That is, the inhibition prevents production of CI, which would otherwise occur as a result of increase in the number of the viral genome, due to replication soon after the commitment to lysis post-infection or during prophage induction. Hence, 1A_Lyt_Lys is a quasi-minimal, not minimal, model. The *pRM* promoter possesses very weak basal expression, whereas *lys* in 1A_Lyt_Lys (and also 1B_Lyt_Lys) lacks basal expression and is activated by Lyt. This difference can be neglected because transcription from CII-activated *pRE* is much higher than basal expression of *pRM*. Schmeissner et al. showed that transcription from CII-activated *pRE* promoter was roughly 3 times the basal expression of the *pR* promoter [18], which is already a much stronger promoter than *pRM*.

### 2.3. Variants of 1A_Lyt_Lys and the mutual repression motif

In order to further investigate design principles of the lysis/lysogeny switch, the mutual repression motif (called here 2_Lyt_Lys) was included because this motif is shown to be able to produce the lysis/lysogeny switching behaviour, at least in deterministic simulation [7], and the core of the GRN responsible for lambda’s lysis/lysogeny decision includes a mutual repression, formed by Cro repression of the *pRM* promoter and CI repression of the *pR* promoter.

#### 2_Lyt_Lys_(Mutual repression)

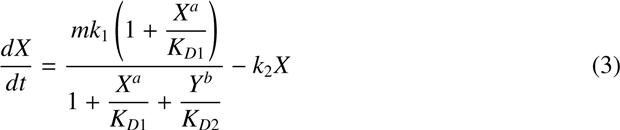

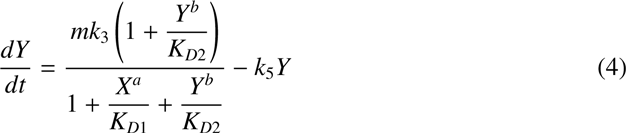

1A_Lyt_Lys and 2_Lyt_Lys have two features common between them, namely constitutive expression of *lyt* and its inhibition by Lys. The presence of these two features can be justified from first principles as follows. Because a single phage infection predominantly causes lysis, where Lyt dominates over Lys, expression of *lyt* has to be strongly constitutive and *lys* should have weak basal expression, if at all. Further, because the infection by multiple phages results in lysogeny, where Lys dominates over Lyt, expression of *lyt* must be suppressed by Lys. We constructed four additional two-protein models by modifying 1A_Lyt_Lys and 2_Lyt_Lys in a way that the resulting models might still be able to produce the switch. The additional models also possessed the two features.

#### 1B_Lyt_Lys

It was suspected that in 1A_Lyt_Lys Lyt self-repression at MoI of 2 could be dispensable. That’s because, firstly, Lys performs the same function much more strongly than Lyt by virtue of being present at a much higher concentration than the latter, whose production is repressed by Lys at this MoI. Secondly, in a negative autoregulatory loop, the protein equilibrium level generally increases with the gene copy number. Under the condition of very strong repression, the protein equilibrium concentration remains nearly the same, but never reduces, with increasing the gene dosage (Supplemental Information). Hence, Lyt_2_ cannot be even less than Lyt_1_ if the negative feedback is the only regulatory mechanism present. However, removal of Lyt self-repression will increase Lyt equilibrium level at MoI of 1. This effect can be nullified by altering the kinetic parameters associated with transcription of *lyt*, translation and degradation of *lyt* mRNA, and degradation of Lyt in a way that the equilibrium level of Lyt at MoI of 1 remains as before.

Lyt negative autoregulation represents two processes in lambda, namely Cro repression of *pR* and CII repression of Q’s production. Given that the present study takes into account equilibrium concentrations of proteins; arguably, kinetic details of Cro repression of genes regulated by *pR* may be ignored. Therefore, arguments given to justify removal of Lyt self-repression can also be employed to eliminate Cro repression of *pR*. Thus, because removal of Cro repression of *pR* increases the equilibrium concentration of *pR*-regulated proteins at MoI of 1, the repression may be done away with while ensuring that the equilibrium concentration of the proteins remain the same at the MoI. This can be achieved by altering the kinetic parameters associated with transcription from *pR*, translation and degradation of the resulting mRNAs, and degradation of proteins produced from those mRNAs. Cro repression of *pR* at MoI of 2 is dispensable because CI performs the same function much more strongly than Cro, because the former is present at much higher concentrations than the latter and the repression cannot bring down the equilibrium level of *pR*-regulated proteins lower than even their equilibrium level at MoI of 1. Cro repression of *pR* (and also *pL*) likely contributes to bistability at MoI of 2, but even this function of Cro can be subsumed into Cro repression of *pRM*, as explained below in the section *Bistability at MoI of 2 and lytic development’s stability*.

It may be argued that CII inhibition of Q’s production, too, can be done away with; that is, by subsuming it into Lys inhibition of *lyt*. CII inhibition of Q’s production is required for the commitment to lysogeny, which in the present study is constrained to occur at MoI of 2. As previously, if kinetic details of CII inhibition of Q’s production are ignored, the process can be subsumed into repression of *pR* (promoter of *lyt*) by CI (Lys), whose production is triggered by CII, in the following way. CII activates transcription from *pRE*, leading to production of CI, which represses *pR*. This results into the repression of *Q*, as its transcription is under the control of *pR*. Thus, CII −| Q is replaced by CII → CI (Lys) −|*pR* → Q (Lyt). This assumption becomes stronger in the light of the experimental result that, as compared to CI, CII has very low impact in reducing production of Q [3]. In a lambda strain lacking the *cI* gene, it was demonstrated that the level of Q protein, reported by its anti-termination of transcription from pR’-tR’-*gfp* promoter-terminator fusion, shows almost no change as a function of MoI (see figure 3D of ref. 3).

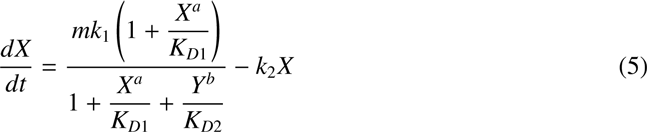

**Figure 3:**
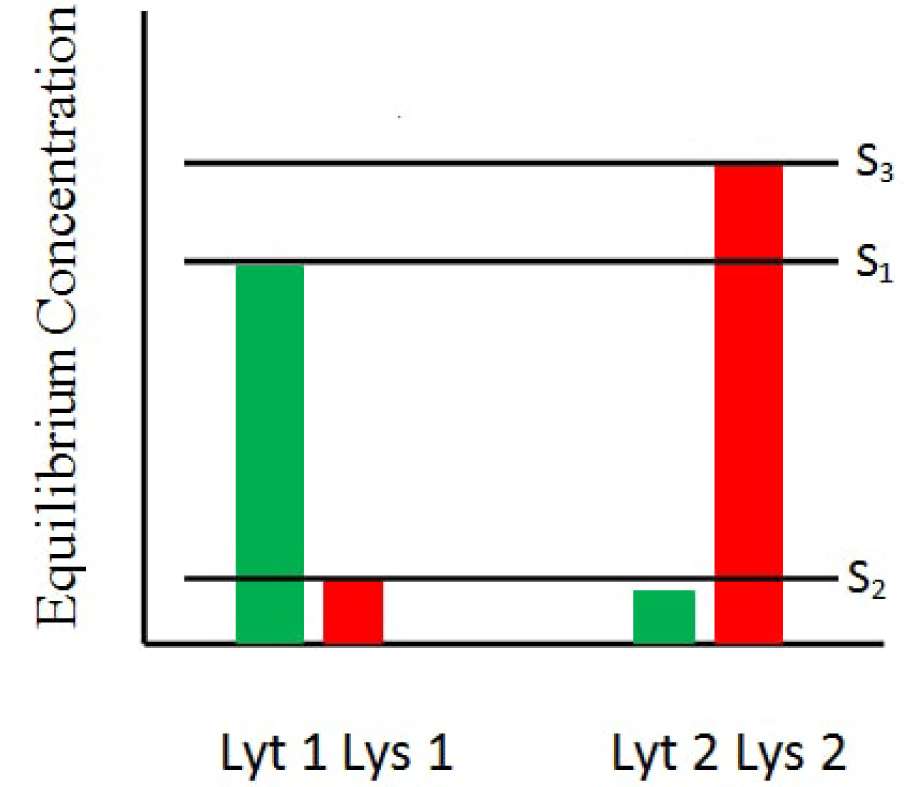
Schematic of the switching behaviour. S_1_ = min{Lyt_1_, Lys_2_}, S_2_ = max{Lys_1_, Lyt_2_}, and S_3_ = max{Lyt_1_, Lys_2_}

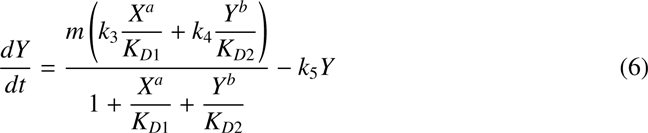

#### 3_Lyt_Lys

This model is a modification of 1B_Lyt_Lys, in that Lyt activation of promoter of *lys* is replaced by basal expression of *lys*. Thus, the basal expression of *lys*, not Lyt activation of *lys* expression, triggers positive autoregulatory loop of Lys in this model.

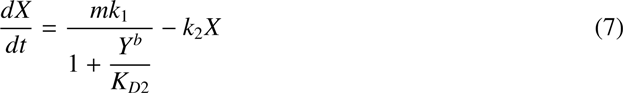

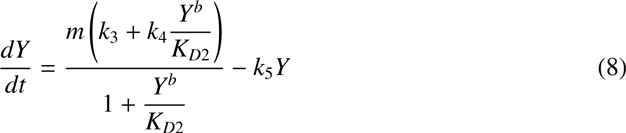

#### 4_Lyt_Lys

Because absolutely essential features of the two-protein models are basal expression of *lyt* and Lys repression of *lyt*, it was be suspected that a model simpler than 2_Lyt_Lys could produce the switch. That is, one could get rid of at least one interaction in 2_Lyt_Lys without rendering its switching behaviour dysfunctional. Thus, one may speculate that Lyt inhibition of *lys* expression in 2_Lyt_Lys may be dispense with, because Lyt is already absent (or at most present at low levels) at MoI of 2 and the lack of Lyt inhibition of *lys* expression might, at least partially, be compensated by reduction in the basal expression of *lys*. Thus we arrive at 4_Lyt_Lys, which comprises of a basal expression of *lys* in addition to two features common in every model.

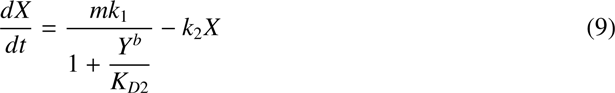

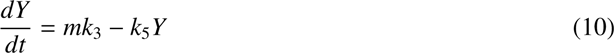

The last four models, which lack Lyt self-repression, namely 1B_Lyt_Lys, 2_Lyt_Lys, 3_Lyt_Lys, and 4_Lyt_Lys, can be placed in a table based on the following three conditions (in addition to two features common in every model): a) whether *lys* possesses basal expression or is activated by Lyt for the purpose of triggering positive autoregulatory loop of Lys (3_Lyt_Lys vs. 1A_Lyt_Lys and 1B_Lyt_Lys), b) whether Lys positive autoregulation is present or not (3_Lyt_Lys vs. 4_Lyt_Lys), and c) whether Lyt inhibition of *lys* expression, ultimately forming the mutual repression motif, is present or not (2_Lyt_Lys vs. 4_Lyt_Lys). Because Lyt inhibition of *lys* expression can be present only when *lys* possesses basal expression, there are four models when *lys* has basal expression; and where its expression is activated by Lyt, there are two models (Table 1). The two additional models are described below.

**Table 1:** Classification of additional two-protein models.

| Model | Basal expression of *lys*/<br>Activation of *lys* by *Lyt* | Activation of *lys*<br>by *Lys* | Inhibition of *lys*<br>by *Lyt* |
| --- | --- | --- | --- |
| 1B_Lyt_Lys | Activation | Yes | N/A |
| 5_Lyt_Lys | Activation | No | N/A |
| 3_Lyt_Lys | Basal | Yes | No |
| 6_Lyt_Lys | Basal | Yes | Yes |
| 4_Lyt_Lys | Basal | No | No |
| 2_Lyt_Lys | Basal | No | Yes |

#### 5_Lyt_Lys

It consists of Lyt activation of *lys* expression in addition to two common features.

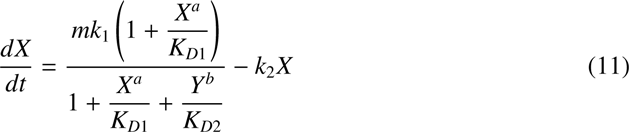

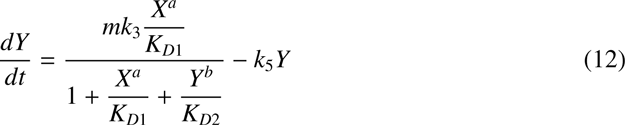

#### 6_Lyt_Lys

*lys* has a basal expression, is activated by Lys, and inhibited by Lyt in addition to two common features. Notably, adding Lyt self-repression to this model makes it identical to the two-protein model of Weitz et al. [6].

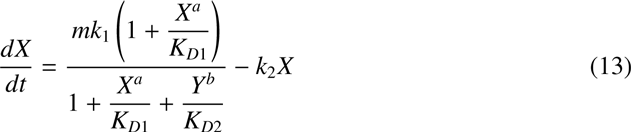

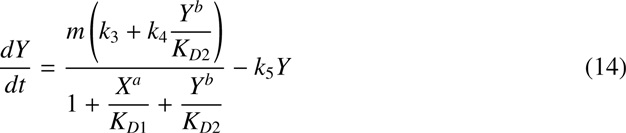

### 2.4. Deterministic simulation

The models were evaluated with respect to the quality of the switch (named here: switch quotient) they generated upon solving their defining equations deterministically for a given set of the rate constants and equilibrium dissociation constants. Because Cro and CI bind as dimer and tetramer, respectively, the Hill coefficient for Lyt and Lys binding are taken to be 2 and 4, respectively. However, for the sake of completeness we also considered Lys binding as a dimer. The rate constants and equilibrium dissociation constants were searched (see Methods) in two stages: the order search and linear search (as they are called here). Only those parameter sets were considered that gave positive switch quotients. Switch quotient corresponding to parameter set generated in deterministic simulations is named deterministic switch quotient (DSQ).

The switch quotient was defined as (See Methods).

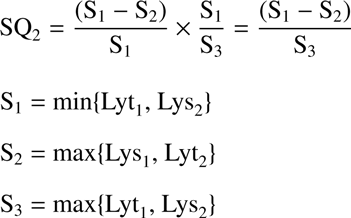

Every DSQ turned out to be finite. As Table 2 shows, all of the models possessing Lys positive autoregulation had the average DSQ of more than 0.97 for both Hill coefficient sets (lowest DSQ among all of the models in this category was 0.9270). The mutual repression motif for the Hill coefficient set a=2, b=2 has only one parameter set, whose DSQ is 0.6666; and for the set a=2, b=4, the average DSQ was 0.9746 (the lowest DSQ was 0.9362). 4_Lyt_Lys for the Hill coefficient set a=2, b=2 gives DSQs of 0.4794 and 0.4707; and for the set a=2, b=4, both DSQs were almost 0.5. DSQs of 4_Lyt_Lys being around and less than 0.5 can be explained in the following way. Because of the absence of Lyt repression of promoter of *lys*, the increase in the genome copy number for this model leads to proportional increase in the equilibrium activity of promoter of *lys*. This causes Lys_1_ to be half of Lys_2_. Hence, according to the definition of the switch, DSQ can be at most 0.5.

**Table 2:** The average and standard deviation of deterministic and stochastic switch quotients for a given range of stochastic success rate for time length of 100 arb. units

| Model | AVG Deterministic SQ<br>(SD) |  | AVG Deterministic SQ<br>(SD)<br>(SSR<sup>a</sup> ≥ 95) |  | AVG Stochastic SQ<br>(SD)<br>(SSR ≥ 95) |  |
| --- | --- | --- | --- | --- | --- | --- |
|  | a=2, b=2 | a=2, b=4 | a=2, b=2 | a=2, b=4 | a=2, b=2 | a=2, b=4 |
| 1A_Lyt_Lys | 0.9917<br>(0.0106) | 0.9896<br>(0.0049) | 0.9898<br>(N/A) | 0.9905<br>(0.0043) | 0.7997<br>(N/A) | 0.6204<br>(0.0624) |
| 1A_Lyt(1)_Lys<sup>b</sup> | 0.9950<br>(0.0053) | 0.9923<br>(0.0045) | none | none | none | none |
| 1B_Lyt_Lys | 0.9806<br>(0.0236) | 0.9769<br>(0.0277) | 0.9971<br>(N/A) | 0.9270<br>(N/A) | 0.7995<br>(N/A) | 0.7679<br>(N/A) |
| 1B_Lyt(1)_Lys | 0.9780<br>(0.0252) | 0.9869<br>(0.0199) | none | 0.9304<br>(N/A) | none | 0.7520<br>(N/A) |
| 2_Lyt_Lys | 0.6666<br>(N/A) | 0.9746<br>(0.0248) | none | none | none | none |
| 3_Lyt_Lys | 0.9950<br>(0.0073) | 0.9893<br>(0.0150) | none | none | none | none |
| 4_Lyt_Lys | 0.4751<br>(0.0043) | 0.4956<br>(0.0006) | none | 0.4956<br>(0.0006) | none | 0.2983<br>(0.0102) |
| 6_Lyt_Lys | 0.9984<br>(N/A) | 0.9845<br>(0.0166) | none | none | none | none |
| Lyt_Lys_CII | 0.9855<br>(0.0155) | N/A | 0.9573<br>(N/A) | N/A | 0.7526<br>(N/A) | N/A |
| Lyt_Lys_CII(1) | 0.9801<br>(0.0151) | N/A | 0.9718<br>(N/A) | N/A | 0.7595<br>(N/A) | N/A |
| Lyt(1)_Lys_CII(1) | 0.9894<br>(0.0134) | N/A | none | N/A | none | N/A |
| Lyt_Lys_CII'<sup>c</sup> | 0.9840<br>(0.0126) | N/A | none | N/A | none | N/A |

<sup>a</sup> SSR = Stochastic success rate
<sup>b</sup> Value in the parenthesis in a model name is the Hill coefficient of protein binding
<sup>c</sup> Parameter sets selected for bistability at MoI of 2.
N/A is used for cases with only one data point.

5_Lyt_Lys did not generate any parameter set. That’s because, in the absence of Lys positive autoregulation, *lyt* needs to have strong basal expression in order to sustain high levels of Lys, whose gene is activated by Lyt, at MoI of 2. That is, along with Lys, Lyt also is expressed at high concentrations at MoI of 2. Equivalently, high Lyt concentration at MoI of 1 also leads to excessive production of Lys at the same MoI. Thus, the two proteins are present in similar amounts at both MoIs. This can be looked in an alternative way. Qualitatively, 5_Lyt_Lys possesses an overall negative autoregulation of Lyt. As argued above, this means that Lyt_2_ cannot be less than Lyt_1_. Thus if Lyt is present at a high concentration at MoI of 1, it’s concentration is even higher at MoI of 2.

In order to examine the significance of cooperativity in Lys positive autoregulation here, the Hill coefficient set a=2, b=1 was considered for every model that possessed the positive feedback. Highest DSQ for 1A_Lyt_Lys turned out to be 0.1262; for 1B_Lyt_Lys: 0.1150; 3_Lyt_Lys: 0.0001; 6_Lyt_Lys: ≈ 0. Extremely low values of highest DSQs generated by this set of Hill coefficients underscores the absolute necessity of cooperative binding of Lys.

### 2.5. Closer to the lambda’s GRN: the minimal three-protein model

We chose to study a three-protein simplified version of the lambda’s switch formed by adding a CII-like protein to 1A_Lyt_Lys, because several studies indicated that CII plays a crucial role in the lysis/lysogeny decision [1,3,6,8]. Besides the addition of a CII-like protein, role of Lyt was modified. That is, in addition to repressing transcription of its own gene and, as a result, the gene co-transcribed with *lyt*, namely *cII*, now Lyt represses transcription of *lys* as well. Thus, roles of Lyt in this model are identical to those of Cro in the lambda’s GRN. As in the two-protein models, functions of Lys here are identical to the functions of CI in the lambda’s GRN. The role of CII in the three-protein model is to activate transcription of *lys*. This corresponds to CII activation of the *pRE* promoter of the lambda’s GRN, resulting in CI production. Like in 1A_Lyt_Lys, CII inhibition of Q’s production is subsumed into Lys inhibition of transcription of *lyt*-*cII*. The three-protein model considered here is different from that in [6], in which CII activates transcription of *cI* from a distinct promoter, *pRE*. In the three-protein model considered here, CII has to compete with Lyt, which represses transcription of *lys*, for binding to the intergenic region and consequently activating promoter of *lys*.

The model equations for the three-protein model are as follows.

Transcription of *lyt*-*cII* genes:

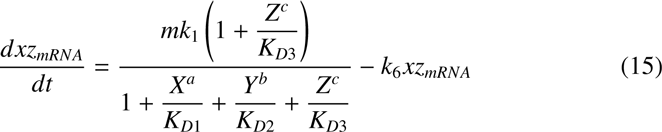

Translation of *lyt*:

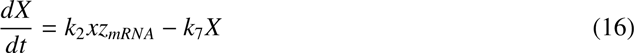

Translation of *cII*:

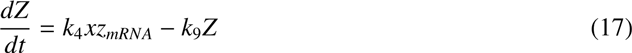

Production of Lys:

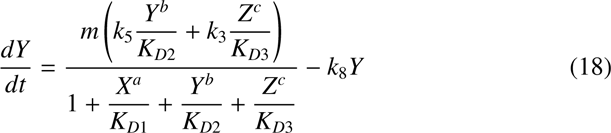

where, c is the Hill coefficient of CII binding; *k*_1_ is the basal rate of transcription of *lyt*-*cII* mRNA; *k*_6_ is the degradation rate of *lyt*-*cII* mRNA; *K_D_*_3_ is the effective equilibrium dissociation constant of CII (see Methods); *k*_2_ and *k*_4_ are the translation rates of *lyt* and *cII*, respectively. *k*_5_ and *k*_3_ are the rate constants for transcriptional activation of *lys* by Lys and CII, respectively. The degradation rate of *lyt*-*cII* mRNA (*xz_mRNA_*), namely *k*_6_, was arbitrarily chosen to be unity to keep the computation time low. The degradation rate of Lyt (*X*), CII (*Z*), and Lys (*Y*), namely *k*_7_, *k*_9_, *k*_8_, respectively, were taken to be unity for the same reason that the degradation rate in the two-protein models were set equal to 1. Because for 1A_Lyt_Lys DSQs generated by the Hill coefficient set a=2, b=2 were as high as DSQs generated by a=2, b=4, applying occam’s razor, the Hill coefficient for Lyt and Lys binding in the three-protein model was taken to be 2 and 2, respectively, and not 2 and 4. Further, taking lead from here, the Hill coefficient of CII binding was taken to be 2, even though it has been shown to exist as tetramer in solution [21] and in crystallized free and DNA-bound state [31, 22].

### 2.6. Stochastic simulation

Because gene expression is stochastic [23, 24], it was crucial to verify if the protein networks that produced the switching behaviour deterministically were also able to function accurately in the presence of noise. Therefore, stochastic simulations were performed by implementing Gillespie algorithm [25], using parameter sets obtained from the deterministic simulations. Barring a few parameter sets of two models for the simulation time length of 200 arb. units, no parameter set was able to produce the switching behaviour in every run of the stochastic simulations. That was because either the SSQ was negative (S_1_ < S_2_) or, rarely, S_3_ was zero. The percentage of runs that produced finite and positive SQ during stochastic simulation for a given parameter set and the Hill coefficient set would henceforth be referred to as stochastic success rate (SSR).

For any given parameter set of any of the models, SSQ was less than its deterministic counterpart (DSQ). As Table 2 and 3, respectively, show for SSR ≥ 90 and 95 >SSR ≥ 90, the average SSQ for the Hill coefficient set a=2, b=2 was greater than that for the Hill coefficient set a=2, b=4 for all of the models. For all of the models and for either of the ranges of SSR, the lowest SSQ for the former Hill coefficient set was greater than the highest SSQ for the latter Hill coefficient set. As Table 5 shows, maximum SSR across all of the models, barring a few cases, increases with the time length of simulation. Also, the order of models arranged according to their maximum SSR remains more or less the same across all of the four time lengths of the simulations. For a given model, for those time lengths which generated at least one parameter set with SSR being within at least one of the two intervals, DSQs (and their average) and SSQs (and their average) were similar to those of the time length of 100 arb. units for the respective intervals of SSR (see Supplementary information Table 1-6). 4_Lyt_Lys has highest maximum SSR, closely followed by 1B_Lyt(1)_Lys, 1B_Lyt_Lys, 1A_Lyt_Lys, and 1A_Lyt(1)_Lys, in that order. Maximum SSR of 3_Lyt_Lys is around halfway between that of 1B Lyt(1) Lys and 6_Lyt_Lys, which has second lowest maximum SSR. 2_Lyt_Lys has lowest maximum SSR. As Table 4 shows, maximum SSR of 6_Lyt_Lys and 2_Lyt_Lys are lower than 80. These two models possess double-negative feedback, or the mutual repression motif. This result that lambda-like switches based on the mutual repression motif work poorly in the presence of noise has been reported earlier also. Avlund et al. showed that various two-protein minimal models, vast majority of which were based on the mutual repression motif, of the lambda’s switch were able to carry out the deterministic counting task, but did not function when noise was introduced [8]. Notably, one of their two two-protein models (i.e., motif b of figure 2 of Avlund et al. [8]) which could pass the counting task even in the presence of noise (though with much lower success as compared to their three-protein models) is 6_Lyt_Lys in the present paper.

**Table 3:** The average and standard deviation of deterministic and stochastic switch quotients for a range of stochastic success rate for time length of 100 arb. units

| Model | AVG Deterministic SQ<br>(SD)<br>(95 >SSR<sup>a</sup> ≥ 90) |  | AVG Stochastic SQ<br>(SD)<br>(95 >SSR ≥ 90) |  |
| --- | --- | --- | --- | --- |
|  | a=2, b=2 | a=2, b=4 | a=2, b=2 | a=2, b=4 |
| 1A_Lyt_Lys | 0.9937<br>(0.0025) | 0.9883<br>(0.0068) | 0.8087<br>(0.0382) | 0.5728<br>(0.1003) |
| 1A_Lyt(1)_Lys<sup>b</sup> | 0.9945<br>(N/A) | 0.9930<br>(N/A) | 0.8084<br>(N/A) | 0.5825<br>(N/A) |
| 1B_Lyt_Lys | 0.9950<br>(N/A) | 0.9972<br>(0.0020) | 0.7891<br>(N/A) | 0.6691<br>(0.0855) |
| 1B_Lyt(1)_Lys | none | none | none | none |
| 2_Lyt_Lys | none | none | none | none |
| 3_Lyt_Lys | none | none | none | none |
| 4_Lyt_Lys | none | none | none | none |
| 6_Lyt_Lys | none | none | none | none |
| Lyt_Lys_CII | 0.9808<br>(0.0008) | N/A | 0.7614<br>(0.0211) | N/A |
| Lyt_Lys_CII(1) | 0.9593<br>(N/A) | N/A | 0.7551<br>(N/A) | N/A |
| Lyt(1)_Lys_CII(1) | 0.9647<br>(N/A) | N/A | 0.7178<br>(N/A) | N/A |
| Lyt_Lys_CII'<sup>c</sup> | 0.9791<br>(0.0106) | N/A | 0.6705<br>(0.0466) | N/A |

<sup>a</sup> SSR = Stochastic Success Rate
<sup>b</sup> Value in the parenthesis is the Hill coefficient of protein binding
<sup>c</sup> Parameter sets selected for bistability at MoI of 2.
N/A is used for cases with only one data point.

**Table 4:** Number of parameter sets for various ranges of stochastic success rate.

| Model | SSR<sup>a</sup> ≥ 95 |  | 95 > SSR ≥ 90 |  | 90 > SSR ≥ 80 |  | Total no. of parameter sets |  |
| --- | --- | --- | --- | --- | --- | --- | --- | --- |
|  | a=2 | a=2 | a=2 | a=2 | a=2 | a=2 | a=2 | a=2 |
|  | b=2 | b=4 | b=2 | b=4 | b=2 | b=4 | b=2 | b=4 |
| 1A_Lyt_Lys | 1 | 4 | 5 | 3 | 10 | 3 | 20 | 17 |
| 1A_Lyt(1)_Lys | 0 | 0 | 1 | 1 | 3 | 9 | 17 | 15 |
| 1B_Lyt_Lys | 1 | 1 | 1 | 5 | 2 | 2 | 6 | 11 |
| 1B_Lyt(1)_Lys | 0 | 1 | 0 | 0 | 8 | 6 | 14 | 10 |
| 2_Lyt_Lys | 0 | 0 | 0 | 0 | 0 | 0 | 1 | 6 |
| 3_Lyt_Lys | 0 | 0 | 0 | 0 | 1 | 6 | 10 | 12 |
| 4_Lyt_Lys | 0 | 2 | 0 | 0 | 0 | 0 | 2 | 2 |
| 6_Lyt_Lys | 0 | 0 | 0 | 0 | 0 | 0 | 1 | 13 |
| Lyt_Lys_CII | 1 | N/A | 2 | N/A | 1 | N/A | 9 | N/A |
| Lyt_Lys_CII(1) | 1 | N/A | 1 | N/A | 5 | N/A | 9 | N/A |
| Lyt(1)_Lys_CII(1) | 0 | N/A | 1 | N/A | 1 | N/A | 9 | N/A |
| Lyt_Lys_CII' | 0 | N/A | 2 | N/A | 4 | N/A | 9 | N/A |

<sup>a</sup> SSR = Stochastic Success Rate

**Table 5:** Highest stochastic success rate for the four time lengths

| Model | Highest stochastic success rate |  |  |  |
| --- | --- | --- | --- | --- |
|  | T = 25 | T = 50 | T = 100 | T = 200 |
| 1A_Lyt_Lys | 89.4 | 93.6 | 96.8 | 100 |
| 1A_Lyt(1)_Lys | 83.8 | 84 | 91 | 97.2 |
| 1B_Lyt_Lys | 90.8 | 93.4 | 97.2 | 99.2 |
| 1B_Lyt(1)_Lys | 96 | 96.6 | 99.2 | 100 |
| 2_Lyt_Lys | 39.4 | 40.2 | 46.4 | 50.6 |
| 3_Lyt_Lys | 74.8 | 81.2 | 86 | 93.8 |
| 4_Lyt_Lys | 87.4 | 93 | 98.8 | 99.8 |
| 6_Lyt_Lys | 57 | 64.8 | 71.2 | 83.8 |
| Lyt_Lys_CII | 92.5 | 94.5 | 95 | 96.5 |
| Lyt_Lys_CII(1) | 90 | 96 | 95 | 97 |
| Lyt(1)_Lys_CII(1) | 88 | 87 | 94.5 | 97.5 |
| Lyt_Lys_CII'<sup>a</sup> | 83 | 85.5 | 92.5 | 96.5 |
| Lyt_Lys_CII(1)'<sup>a</sup> | 32.5 | 35 | 39 | 43 |

<sup>a</sup> Parameter sets selected for bistability at MoI of 2.

### 2.7. Bistability at MoI of 1 and lysogen stability

After the commitment to lysogeny (at MoI of 2 in the present study), if only one of the phage genomes gets integrated into the bacterial chromosome, thereby bringing the MoI down to 1, maintenance of lysogeny will not be possible and lysis will ensue, in case only one stable state existed at MoI of 1. That’s because a single phage will not be able to sustain high equilibrium level of CI, produced by two phages. In deterministic simulations, every two-protein model that possessed Lys positive autoregulation exhibited bistability at MoI of 1 for every parameter set, except one that belonged to 1B_Lyt_Lys. In the other stable state, Lyt concentration is almost zero and that of Lys is about half of its concentration at MoI of 2.

For 4_Lyt_Lys, none of the parameter sets produced bistability at MoI of 1. That’s because the nullcline of Lys is a flat line and that of Lyt is a monotonically decreasing function. Therefore, 4_Lyt_Lys cannot produce bistability at either of the MoIs. For 2_Lyt_Lys, only one parameter set generated bistability at MoI of 1 for the Hill coefficient set a=2, b=2. For the Hill coefficient set a=2, b=4, all parameter sets except one exhibited bistability at MoI of 1. It turned out that the parameter set which did not produce bistability has *K_D_*_1_ and *K_D_*_2_ about 5 orders of magnitude higher than the average *K_D_*_1_ and *K_D_*_2_, respectively, of the rest of the parameter sets of 2_Lyt_Lys (Supplementary Information: Table 10). Because in 4_Lyt_Lys Lyt does not bind to any promoter, it can be thought to be having an extremely large equilibrium dissociation constant (i.e., very large *K_D_*_1_). Further, *K_D_*_2_ of the parameter set is within an order of the average *K_D_*_2_ of parameter sets of 4_Lyt_Lys for the given set of Hill coefficients. Therefore, it can be argued that the parameter set actually belonged to 4_Lyt_Lys.

Every three-protein model for every parameter set exhibited bistability at MoI of 1. The values of Lyt and Lys of second stable state at MoI of 1 in the three-protein models were about the same as the values of Lyt and Lys, respectively, of second stable state in the two-protein models at MoI of 1. Stochastic bistability was searched using the same method employed to search deterministic bistability. However, stochastic bistability could not be detected for any of the two-protein models and the three-protein model at MoI of 1. This may be due to noise driving trajectories out of the basin of attraction of one stable state to that of the other [26]. Stochastic bistability was checked only for those parameter sets whose SSR was more than 95%. This was done to avoid mistaking a failure to generate the switching behaviour for bistability.

A study showed that CI positive autoregulation is dispensable if basal expression of the *pRM* promoter is enhanced [30]. Removing Lys positive autoregulation from the three-protein minimal model and conferring basal expression to *lys* transforms the model to a mutual repression motif with two additional features, namely CII activation of *lys* expression (which represents CII activation of *pRM*) and Lyt inhibition of promoter of *lyt*-*cII* (which represents Cro inhibition of *pR*). The models which have exhibited bistability at MoI of 1 at least either had Lys self-activation or mutual repression between Lyt and Lys. Removing Lyt inhibition of *lys* expression transforms 2_Lyt_Lys to 4_Lyt_Lys, which cannot have bistability at all MoIs. Hence, because eliminating Lyt repression of *lys* expression in 2_Lyt_Lys makes the system monostable at all MoIs, a testable prediction of the present study is that removing Cro repression of *pRM* in the modified phage of the study should render it unable to cause lysogeny, which requires bistability at MoI of 1.

### 2.8. Bistability at MoI of 2 and lytic development’s stability

Because replication of viral genome after the commitment to lysis causes the MoI to become more than 1 (2 in the present study), lytic development will get affected and lysogeny might ensue, in case only one stable state existed at MoI of 2. The consequence of lack of bistability at MoI of 2 is not identically opposite to that of lack of bistability at MoI of 1. While without bistability at MoI of 1 lysogeny is reversed in the simulations (though to the author’s knowledge, experimental observation of this is lacking), lack of bistability at MoI of 2, or higher MoIs in general, could still lead to lysis and with appreciable burst size in the real system. This is because once Q protein activates the lytic genes, the commitment to lysis occurs and CI is not able to affect late lytical processes. However, CI can still affect lytical processes of the genomes which are produced as a result of the ongoing replication, more so because the presence of larger number of genomes leads to greater production of CI. The lack of bistability at higher MoIs impacts prophage induction vastly more than lytic development post-infection [27]. This is discussed later in this section.

In deterministic simulations, only two models, namely 2_Lyt_Lys and 6_Lyt_Lys, exhibited bistability at MoI of 2. Notably, among all of the two-protein models, only these two models possess Lyt repression of *lys* expression. In the other stable state, Lyt concentration is double of its concentration at MoI of 1, and that of Lys is almost zero. For 2_Lyt_Lys, the parameter set that produced bistability at MoI of 1 for the Hill coefficient set a=2, b=2 is the only parameter set that exhibited bistability at MoI of 2. For the Hill coefficient set a=2, b=4, all parameter sets (except the one which actually belonged to 4_Lyt_Lys) exhibited bistability at MoI of 2. For 6_Lyt_Lys, one parameter set generated bistability at MoI of 2 for the Hill coefficient set a=2, b=2. For the Hill coefficient set a=2, b=4, all parameter sets except one produced bistability at MoI of 2.

None of the three-protein models for any parameter set exhibited bistability at MoI of 2. However, given that the three-protein models possess Lyt repression of *pR*, the only feature absolutely associated with bistability at MoI of 2 for the two two-protein models, it was suspected that the three-protein models also could exhibit bistability at MoI of 2 if the property was actively selected for. Therefore, the parameter search was carried out by adding another selective filter. That is, Lyt at MoI of 1 (when the model equations were solved with the initial values of Lyt and Lys being equal to zero) <Lyt at MoI of 2 (when the model equations were solved with the initial values of Lyt and Lys being their equilibrium values at MoI of 1). This method generated 9 parameter sets for Lyt_Lys_CII (named Lyt_Lys_CII**^’^**) and 3 for Lyt_Lys_CII(1) (named Lyt_Lys_CII(1)**^’^**) but none for Lyt(1)_Lys_CII(1) (named Lyt(1)_Lys_CII(1)**^’^**).

Further, while maximum SSR of Lyt_Lys CII**^’^** was very close to that of Lyt_Lys_CII, maximum SSR of Lyt_Lys_CII(1)**^’^** was less than half of that of Lyt_Lys_CII(1). This result underscores the importance of cooperative binding for all of the three key proteins of the switch. The most appreciable change between Lyt_Lys_CII and Lyt_Lys_CII**^’^**) was in the equilibrium dissociation constants. The average *K_D_*_1_ and *K_D_*_2_ of Lyt_Lys_CII**^’^**) were about an order of magnitude lower than those of Lyt_Lys_CII, whereas the average *K_D_*_3_ was about two orders of magnitude lower for the same comparison (Supplementary Information: Table 11). Notably, the parameter search with the additional selective filter did not produce any parameter set (i.e., which could exhibit bistability at MoI of 2) for the two-protein models other than 2_Lyt_Lys and 6_Lyt_Lys. This further confirms that Lyt (or Cro) repression of *lys* (or *pRM*) is required to produce bistability at MoI of 2.

A study demonstrated that the average number of phages produced per infection (burst size) of a mutant defective in Cro repression of *pRM* was 75% of that of the wild type, whereas burst size produced after prophage induction triggered by ultra violet (UV) at 10 J/m^2^ UV and at 5 J/m^2^ UV in mutants were 1/3 and 1/8 times of that in the wild type, respectively [27]. The lack of Cro repression of *pRM* having much larger impact on burst size in prophage induction as compared to infection can be explained in the following way. During UV-triggered induction, the residual CI [29] and CI being produced by residual *cI* mRNAs which remain unaffected by UV irradiation activate multiple *pRM* promoters, lacking Cro repression of *pRM*, and consequently trigger CI positive autoregulation. CI, now present in higher concentrations, represses the *pR* promoter, which regulates the *Q* gene. Thus lytical processes, namely the viral genome replication and the associated protein production, occur to a lesser extent, hence the lower burst size.

On the other hand, for CI to affect lytic development post-infection in phages lacking Cro repression of *pRM*, CI has to be made to the level sufficient to activate *pRM*. But for that to happen, transcription from CII-activated *pRE* and translation of the resulting transcripts has to take place first. This extra time allows more rounds of replication of the viral genome and associated production of viral proteins. Hence the effect of defective Cro repression of *pRM* on burst size during lytic development post-infection is lesser. Although before CI is able to repress *pR*, CII can impede production of Q protein by activating transcription from *paQ*, but, as aforementioned, this process is much weaker than CI repression of *pR* in inhibiting production of Q protein.

It should be noted that the modeling scheme in the present work is unable to embody the fact that even without Lyt repression of *lys* expression, lytic development does occur, albeit to a lesser extent (lower burst size). In other words, in the models which produce the switching behaviour but do not exhibit bistability at MoI of 2, increase in MoI due to genome replication after the commitment to lysis will nullify the commitment and ensuing lytic development (zero burst size), and consequently lysogeny will occur. This inability of the present modeling scheme stems from not considering lytical processes, initiated by Q, explicitly.

The frequency of lysogeny (proportion of infected cells that formed lysogens) post-infection in the mutant was 30% higher than that in the wild type, and the fraction of cells induced at 10 J/m^2^ UV and at 5 J/m^2^ UV was 32% and 48% lower for the same comparisons. The increase in the lysogenization frequency post-infection and decrease in the fraction committing to induction can be explained as being due to defective Cro repression of *pRM* tilting the decision towards lysogenization. However, an additional factor exists for the latter case: activation of *pRM* by residual CI and CI being produced by residual *cI* mRNA. It should be noted that the protein-only approach of the present study cannot take into account CI production by residual *cI* mRNAs during the process of prophage induction.

Cro represses *pL* and *pR* by fourfold and twofold, respectively [28]. Therefore, the absence of Cro increases the level of CII by allowing transcription of *cII*, regulated by *pR*, and *cIII*, regulated by *pL* and whose product prevents degradation of CII by protease HflB. Because CII engenders CI production (by activating *pRE*) and CI activates *pRM*, like Cro repression of *pRM*, Cro inhibition of *pL* and *pR* also should contribute to bistability at MoI of 2. This role of Cro is not modeled in the present study. However, because the end result of CII production is activation of *pRM*; qualitatively, Cro inhibition of *pL* and *pR* could be subsumed into Cro inhibition of *pRM*. As is the case at MoI of 1, stochastic bistability could not be detected for any of the two-protein and three-protein models at MoI of 2. Because stochastic bistability could be checked only for those parameter sets whose SSR was more than 95%, most models, including the two two-protein models which exhibited deterministic bistability at MoI of 2 (i.e., 2_Lyt_Lys and 6_Lyt_Lys), could not be checked for their potential ability to produce stochastic bistability at the MoI of 2.

## 3. Design principles of the lambda’s switch

Because, barring 2_Lyt_Lys for a=2, b=2 and 4_Lyt_Lys for both Hill coefficient set, DSQ generated by every model for every parameter set is more than 0.9, design principles of the lambda’s switch cannot be based on the magnitude of DSQ alone. One noticeable feature that the models clearly differ in is maximum SSR they generate. Also, because bistability at both MoIs is an absolute requirement for the switch, among the two-protein models only 2_Lyt_Lys and 6_Lyt_Lys (2_Lyt_Lys plus self-activation of Lys) qualify as possible switch designs. Of these two, the latter is a better model than the former because of producing higher maximum SSR. 6_Lyt_Lys (3_Lyt_Lys plus Lyt repression of *lys* expression) has lower maximum SSR than 3_Lyt_Lys but higher maximum SSR than 2_Lyt_Lys. Hence, it can intuitively be argued that 6_Lyt_Lys is a compromise between 3_Lyt_Lys (for its high maximum SSR) and 2_Lyt_Lys (for its bistability at MoI of 2). In other words, 6_Lyt_Lys incurs a drop in maximum SSR as compared to 3_Lyt_Lys in lieu of gaining bistability at MoI of 2.

Further, from comparing 1B_Lyt_Lys with 3_Lyt_Lys, it is inferred that upregulation of *lys* expression, in order to trigger positive autoregulatory loop of Lys at MoI of 2, by Lyt binding to promoter of *lys* gives higher maximum SSR than having basal expression of *lys* carry out that function. Applying this cause-effect relationship in an additive fashion to 6_Lyt_Lys, it is posited here that maximum SSR of 6_Lyt_Lys can be improved by replacing basal expression of *lys* with Lyt activation of promoter of *lys*. However, because Lyt represses *lys* in 6_Lyt_Lys, another protein, whose gene is co-transcribed with *lyt*, is employed to activate transcription of *lys* (in a cooperative manner). That multimeric protein is CII [21,31,22]. The three-protein model thus generated is the minimal model for the lambda’s switch. Thus, lambda’s lysis/lysogeny switch is based on three features: a) mutual repression, b) cooperative positive autoregulation of CI, and c) cooperative binding of the activator protein, not basal expression, triggering positive feedback loop formed by CI autoregulation.

A speculative explanation for why 1B_Lyt_Lys has higher maximum SSR than 3_Lyt_Lys is given now. 1B_Lyt_Lys differs from 3_Lyt_Lys with regard to how positive feedback loop of Lys is triggered. In 1B_Lyt_Lys, the loop is triggered by Lyt binding to, and consequently activating, promoter of *lys*, whereas in 3_Lyt_Lys the loop is triggered by basal expression of *lys*. In both models, at MoI of 2, the concentration of Lyt initially becomes more than its equilibrium concentration at MoI of 1 but then comes back to very low level, as shown in the qualitatively representative figure 4A. As figure 4A shows, until Lyt trajectory begins to steeply fall back to low level, the concentration of Lys remains very low; and as the concentration of Lys rises, Lyt level drops. This indicates that the time since beginning of the simulation and until the steep fall of the Lyt trajectory is the period when positive autoregulatory loop of Lys is triggered. Additionally, as also shown in the figure 4A, the concentration of Lys at MoI of 1 remains very low as compared to that of Lyt in the two models. It may be argued that these observations justify approximating Y = 0 in the numerator and denominator of the first term on the right-hand side of equality of equations 4 and 8 for time interval between beginning of simulation and until the point when Lyt trajectory begins to drop sharply. Thus one gets *dY*/*dt* equal to *m*(*k*_3_ *X^a^*/*K_D_*_1_)/(1 + *X^a^*/*K_D_*_1_) – *k*_5_*Y* and *mk*_3_ – *k*_5_*Y*, respectively, – an approximation considered to be valid for the said time interval.

**Figure 4:**
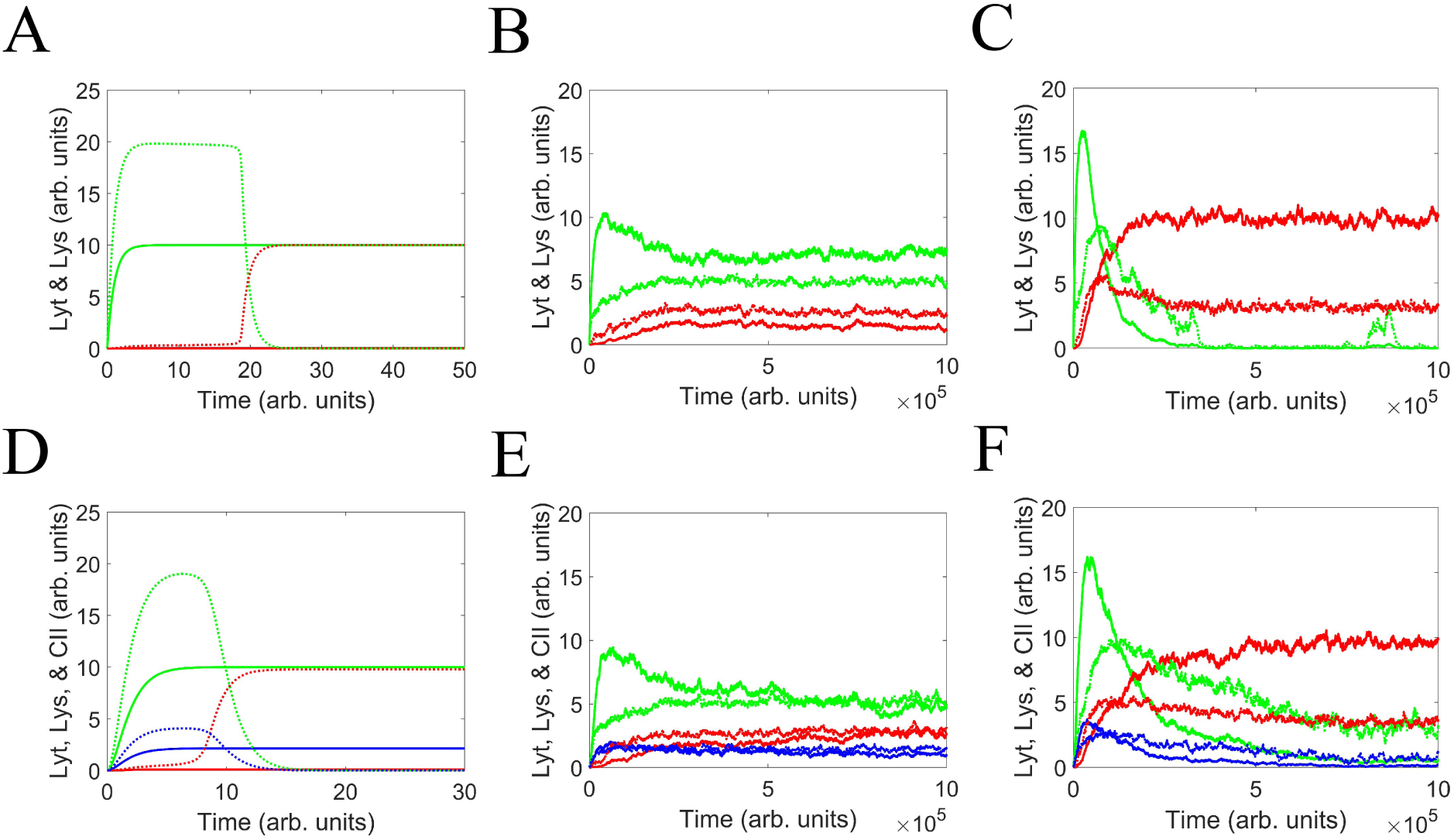
Deterministic and stochastic simulations of the quasi-minimal two-protein model (1A_Lyt_Lys) and the three-protein model (Lyt_Lys_CII). Lyt, Lys, and CII are represented by green, red, and blue, respectively. Protein concentrations at MoI of 1 and 2 in deterministic simulations are depicted by solid curve and dashed curve, respectively. The average and standard deviation of the number of protein molecules as a function of time in stochastic simulations (100 runs) are depicted by solid curve and dotted curve, respectively. For a given model, the parameter set which generated highest stochastic success rate (SSR) for the time length of 100 arb. units was used for these simulations. The averaged stochastic trajectories shown here are qualitatively similar to those of all the other models for parameter sets which generated large values of SSR (data not shown), whereas the deterministic simulation trajectories were so irrespective of the value of SSR (data not shown). In the stochastic simulation plots, the original abscissa, which had unequally-spaced time intervals, was converted to one with equally-spaced time intervals. Each unit of abscissa was divided into 10000 intervals. For the tiny fraction of intervals which still contained more than one event, their last event was defined to be their only event. (**A**) Deterministic simulations of 1A_Lyt_Lys. At MoI of 2, the concentration of Lyt initially becomes more than its equilibrium concentration at MoI of 1 but then comes back to very low levels. The bell-shaped kinetics occurs because the initial rate of Lyt production at MoI of 2 (i.e., when *Y* remains very low) is higher than that at MoI of 1; however, as Lyt concentration rises, positive autoregulatory loop of Lys gets triggered, leading to Lys production in high amount, which represses *lyt*. (**B-C**) Stochastic simulations of 1A_Lyt_Lys for MoI of 1 and 2, respectively. (**D**) Deterministic simulations of Lyt_Lys_CII. At MoI of 2, the concentration of Lyt and CII initially become more than their respective equilibrium concentration at MoI of 1 but later come back to very low levels. A theoretical study of the lysis/lysogeny decision using a three-protein model, which is very similar to that of the present paper, also reported bell-shaped kinetics for CII [6]. Analogous to the two-protein model, the three-protein model displays bell-shaped kinetics because the initial rate of CII production at MoI of 2 (i.e., when *Y* and *Z* remain very low) is higher than that at MoI of 1; however, as CII concentration rises, positive autoregulatory loop of Lys gets triggered, leading to Lys production in high amount, which represses the promoter regulating *lyt* and *cII*. (**E-F**) Stochastic simulations of Lyt_Lys_CII for MoI of 1 and 2, respectively. The bell-shaped curve for CII at MoI of 6 was reported by an experimental study [3]. Curiously, while the equilibrium concentration of CII is higher than that of CI in the deterministic simulation, CII settles down to lower level than CI in the stochastic simulation.

**Figure 5:**
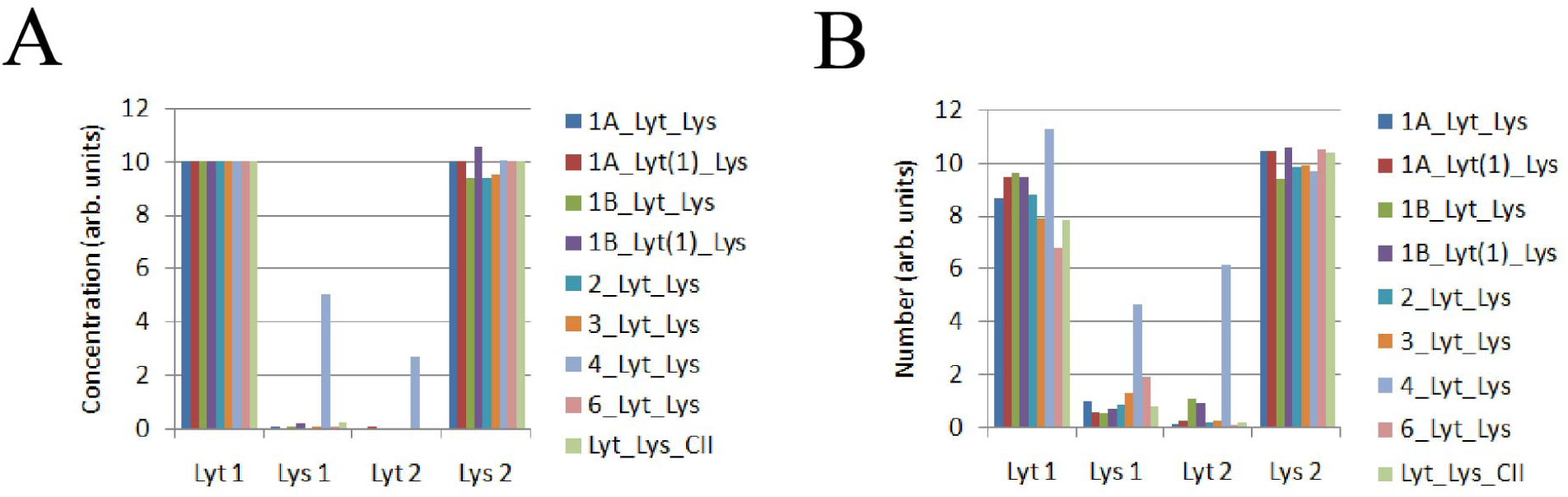
Equilibrium values of Lyt and Lys corresponding to those parameter sets which gave maximum SSR for the time length of 100 arb. units (see Table 5). (**A**) Deterministic simulations. (**B**) Stochastic simulations. Notably, Lys_1_ and Lyt_2_ of 4_Lyt_Lys are much higher than those of the any other model.

**Figure 6:**
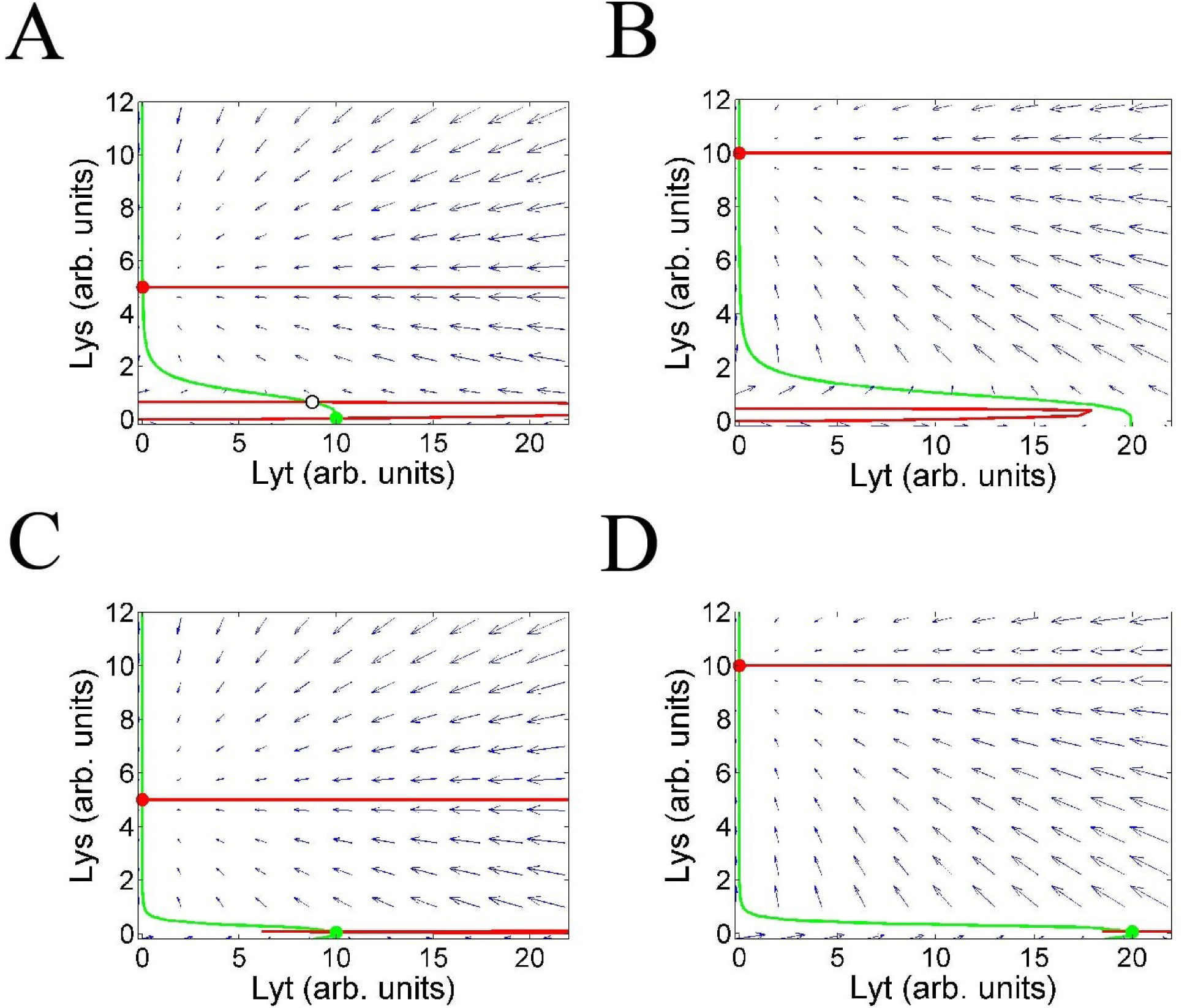
Phase diagrams of 1A_Lyt_Lys (A-B) and 6_Lyt_Lys (C-D) for MoI of 1 and 2 corresponding to their respective parameter set which gave maximum SSR. The green and red full circles are stable fixed points, whereas the empty black circle is unstable fixed point (not marked for 6_Lyt_Lys for either of the MoIs because of being very close to the stable fixed point). (**A**) The green dot shows anti-immune state established by a single phage (MoI of 1), whereas the red dot shows immune state established by two phages (MoI of 2) but which persists even when genome of only one of the phages (MoI of 1) gets integrated into the bacterial chromosome. (**B**) The red dot shows immune state established by two phages (MoI of 2). (**C**) Same as A. (**D**) The red dot shows immune state established by two phages (MoI of 2), whereas the green dot shows anti-immune state established by a single phage (at MoI of 1) but which persists even after the genome replication (MoI of 2)

1B_Lyt_Lys and 3_Lyt_Lys can fail in two ways: either positive autoregulatory loop of Lys gets triggered at MoI of 1 or fails to get triggered at MoI of 2. The first term of the approximated form of *dY*/*dt* for the two models represents triggering of the positive feedback loop, which occurs during the said time interval. For 3_Lyt_Lys, it is equal to *mk*_3_ and, as a result, increases proportionally with MoI. For 1B_Lyt_Lys, the effect of increase in MoI on the first term of the approximated form of *dY*/*dt* depends on the magnitude of the ratio between *X^a^* and *K_D_*_1_. For *X^a^*/*K_D_*_1_ >> 1, the term becomes approximately equal to that for 3_Lyt_Lys. On the other hand, for *X^a^*/*K_D_*_1_ << 1, the term becomes approximately equal to *m*(*k*_3_ *X^a^*/*K_D_*_1_). Because during the said time interval, Lyt concentration at MoI of 2 remains higher than its concentration at MoI of 1, the increase in MoI causes more than proportional increase in the term. This greater separation between the propensity to trigger the autoregulatory loop of Lys at MoI of 1 and 2 for 1B_Lyt_Lys as compared to 3_Lyt_Lys is posited here to be responsible for the former model having higher maximum SSR than the latter. Another, perhaps less likely, reason could be that, while gene expression in 3_Lyt_Lys is ultimately driven by two promoters having independent (basal) expressions, that in 1B_Lyt_Lys is ultimately driven by only one promoter, thereby lending 1B_Lyt_Lys tighter control over overall expression than 3_Lyt_Lys. Because, as argued above, 1B_Lyt_Lys is qualitatively similar to 1A_Lyt_Lys – differing from the latter in only not possessing Lyt self-inhibition and thus having identical equation for the dynamics of Lys – the previous argument also explains why 1A_Lyt_Lys has higher maximum SSR than 3_Lyt_Lys.

## 4. Methods

### 4.1. Derivation of the model equations

In order to calculate the effective equilibrium dissociation constant of proteins, the present work uses the fact that binding of protein to itself or DNA are much quicker processes than transcription and translation. Hence the former processes are assumed to have reached quasi-steady states. In the expressions below, P, X, and Y are promoter, Lys, and Lyt, respectively.

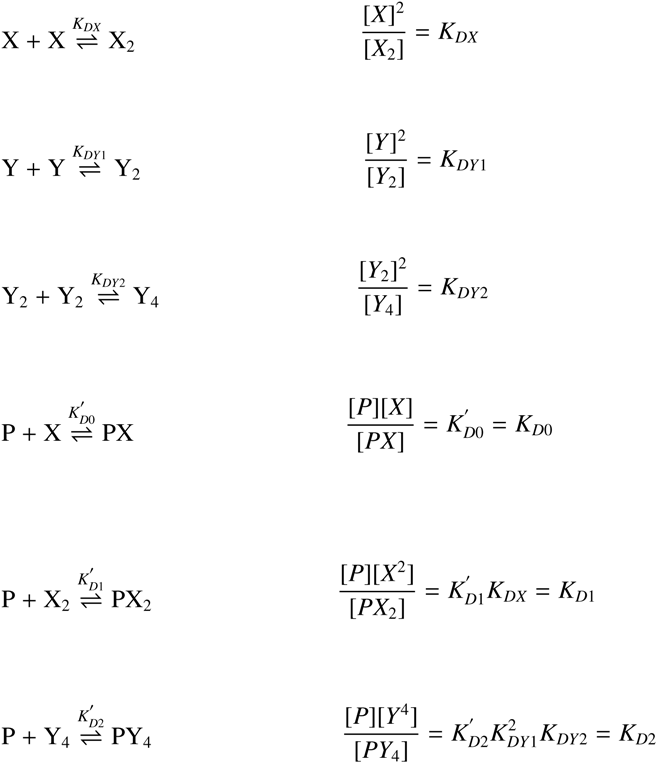

The above expressions for concentrations of promoter-protein complexes are for cases where a) Lyt binds as monomer, b) Lyt binds as dimer, and c) Lys binds as tetramer. They exhaust all other cases, that is, of monomeric and dimeric Lys and CII. Further, transcription and mRNA degradation are assumed to be much faster processes than translation and protein degradation, hence mRNAs also are assumed to have reached quasi-steady states. Because the *lyt* and *cII* genes are under the control of the same promoter, in order to allow for potentially different rates of translation of their corresponding cistrons during the stochastic simulations, the dynamics of *lyt*-*cII* mRNA are considered explicitly. Hence, except that for *lyt*-*CII* mRNA, all of the equations describe the dynamics of only proteins. With the expressions for concentrations of promoter-protein complexes, one can write generalized form of the term representing the rate of protein production.

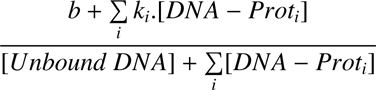

where b, if present, is the basal rate of expression and *k_i_* is the rate constant for transcriptional activation by i*_th_* protein.

### 4.2. Parameter search

Parameter sets, namely the rate constants and equilibrium dissociation constants, of the model equations were searched deterministically in two stages, namely order search and linear search. In the order search, parameters were searched as 3’s exponent, which was varied between −5 and 5 with the difference of 1 (1/243, 1/81, … 1 …, 81, 243), in a nested fashion. Thus the number of parameter sets searched for a given model was equal to 11 raised to the number of parameters. In the interest of saving time, for the three-protein models, the rate of basal expression of *pR* and equilibrium dissociation constant of Lyt and Lys were taken from the parameter set of 1A_Lyt_Lys (for Lyt_Lys_CII and Lyt_Lys_CII(1)) and 1A_Lyt(1)_Lys (for Lyt(1)_Lys_CII(1)) which gave maximum SSR. The model equations were solved deterministically with a given parameter set and the value of DSQ was used to decide whether the parameter set was rejected or not. Also in the interest of saving time, a parameter set was rejected if its DSQ was lower than that of the previously selected parameter set. The vast majority of switch quotients generated by the order search were far from their optimal values because the rate constants and equilibrium dissociation constants were increased geometrically, thereby causing a lot of intervening values to remain unsampled. Therefore, the linear search, which searches the neighbourhood of a parameter set in a linear fashion, was employed in order to further refine parameter sets generated by the order search.

In parameter sets generated by the order search, it was noted that those parameter sets whose DSQs were too close to each other were either rescaled form of each other or differed in those parameters to which DSQ was resilient up to a certain range. Hence, in order to remove the redundancy, the initial parameters for the linear search were taken in such a way that the difference between their consecutive DSQs was at least 0.001. However, because 1A_Lyt_Lys, 1A_Lyt(1)_Lys, 1B_Lyt_Lys, 1B_Lyt(1)_Lys, and all of the three-protein models generated large number of parameter sets, the difference between the consecutive DSQs for these four models was kept to be at least 0.01 in the interest of saving time. This way, Lyt_Lys_CII, Lyt_Lys_CII(1), Lyt(1)_Lys_CII(1) generated 15, 15, and 17 parameters, respectively. However, in order to save time, only their first 9 parameters were considered for the linear search.

The linear search was carried out in the following way. The value of each parameter (say, V) of a set was varied between V-3*V/5 and V+3*V/5 with the increment of V/5, in a nested fashion. That is, a parameter value of, say, 10 was varied between 4 and 16 with the increment of 2. Thus, the number of parameter sets searched in a single round of the linear search was equal to the number of parameters raised to the power 7. However, for the three-protein models, which had eight parameters, in order to save time, each parameter was varied between V-2*V/5 and V+2*V/5 with the increment of V/5, in a nested fashion. A parameter set was rejected if its DSQ was lower than that of the previous parameter set. Thus the linear search was guided by the value of the switch quotient itself. The linear search was ended if the latest DSQ was either lower than the previous one (which never happened) or if ((latest DSQ – previous DSQ)/previous DSQ) was less than 0.01. Again in the interest of saving time, for the three-protein models, the linear search was ended if DSQ at the end of a round was at least 0.95. It should be noted that the linear search is path dependent: a linear search may end up at a local maximum because it treaded a path which yielded higher DSQ initially instead of another path which, in spite of yielding lower DSQ initially, would have reached the global maximum. Optimization could not be used here because the objective function, which had to be based on the ratio of protein equilibrium concentration, was independent of multiplicity of infection, as the latter got cancelled out.

Parameter sets were normalized such that Lyt equilibrium concentration at MoI of 1 becomes equal to 10 arb. units. This was done for two purposes: a) to reduce the possibility of Lyt level at MoI of 1 and Lys level at MoI of 2 dropping to zero in the stochastic simulations; and b) in order to make comparison of parameter sets and protein equilibrium values visually feasible. For both types of searches, simulations were carried out for the time length of 100 arb. units. There was a possibility of a system of equations defining a particular model not reaching the stable state by 100 arb. units for a given parameter set. In order to eliminate such parameter sets, a final simulation was done for the time length of 10^5^ arb. units. Only few parameter sets turned out to not had reached the fixed point, and all of such parameter sets produced negative DSQ.

Avlund et al. [7] use (Lyt_1_/Lys_1_)*(Lys_2_/Lyt_2_) >10 as the criterion for a protein network to be successful in producing the switch. However, this measure cannot distinguish between the following three vastly different scenarios: A) Lyt_1_=100, Lys_1_=1; Lyt_2_=1, Lys_2_=1. B) Lyt_1_=10, Lys_1_=1; Lyt_2_=1, Lys_2_=10. C) Lyt_1_=1, Lys_1_=1; Lyt_2_=1, Lys_2_=100. Initially we considered the difference between minimum of Lyt_1_ and Lys_2_ and maximum of Lys_1_ and Lyt_2_ normalized by the former difference to be the measure of the switch’s quality. This measure is chosen for the following reasons. Arguably, for the switch to be of high quality Lyt_1_ and Lys_2_ should be large and Lys_1_ and Lyt_2_ should be small. Thus, taking the minimalist approach, we considered the normalized difference of the worst of the former two values and the best of the latter two values to be the measure of the switch’s quality.

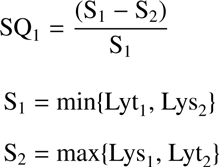

Parameter sets selected by this definition of the switch’ quality usually gave unequal Lyt_1_ and Lys_2_. From the perspective of simplicity, we believe that the two values should be as close as possible. Therefore, the expression is multiplied by the ratio between S_1_ and S_3_ in order to penalize the difference between S_3_ and S_1_. This expression was taken to be the measure of the switch’s quality. This expression, like the previous one, varies between 0 and 1.

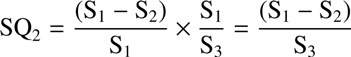

We did not use a measure based on the ratio of protein equilibrium concentration, namely Lyt_1_/Lys_1_ and Lys_2_/Lyt_2_, because of the following reason. Suppose a parameter set gives Lyt_1_ and Lys_1_ equal to 10 arb. units and 1 arb. units, respectively; and Lyt_2_ and Lys_2_ equal to 1 arb. units and 10 arb. units, respectively. Thus the ratios Lyt_1_/Lys_1_ and Lys_2_/Lyt_2_ are both equal to 10. Further, suppose another parameter set gives Lyt_1_ and Lys_1_ equal to 10 arb. units and 1 arb. units, respectively; and Lyt_2_ and Lys_2_ equal to 10 arb. units and 100 arb. units, respectively. The ratios Lyt_1_/Lys_1_ and Lys_2_/Lyt_2_ are both still equal to 10. Although the ratios are the same, the two scenarios are not identical biologically, because if Lyt at the equilibrium concentration of 10 arb. units can cause lysis, there is no reason why it will not do the same when present at the same concentration even in the presence of 100 arb. units, instead of 1 arb. units, of Lys. This is because, although Lys represses Lyt production, Lys has no impact on the imaginary downstream lytic pathway once it is activated by Lyt. In the lambda’s GRN, it translates to CI having no impact on lytic development once the protein Q is made to a sufficient level.

Parameter sets, though not all of them, of only 2_Lyt_Lys and 6_Lyt_Lys exhibited bistability at both MoIs. Therefore for these two models, those parameter sets which did not exhibit bistability at both MoIs were discarded for all purposes. For the models possessing Lys positive autoregulation, those parameter sets which did not exhibit bistability at MoI of 1 were discarded for all purposes (this way only one parameter set, that is, of 1B_Lyt_Lys, was discarded). At MoI of 1, Lyt trajectory exhibits an initial rise, followed by a partial drop, and finally, plateauing. At MoI of 2, the trajectory is bell-shaped: it rises and then finally drops to very low values. The transient kinetics at both MoIs get completed at most by 100 arb. units. In order to calculate stochastic switch quotient (SSQ) and stochastic success rate (SSR), protein levels were averaged for 25, 50, 100, and 200 arb. units of time, keeping the starting time point for averaging at 100 arb. unit. That is, for averaging time=25, the averaging was done between 100 and 125 arb. units; for averaging time=50, the averaging was done between 100 and 150 arb. units; and so forth. All stochastic data, unless specified otherwise, presented here is for time=100 arb. units (for other time lengths, see Supplementary Information: Table 1-9). Only those runs that produced finite and positive SSQ were taken into account in calculating the average and standard deviation of SSQ. For the two-protein and three-protein models, respectively, 500 and 200 simulations were performed for a given parameter set and the Hill coefficient set.

In order to ascertain lysogen stability, the model equations were solved at MoI of 1 with the initial values of Lyt and Lys being their equilibrium values at MoI of 2 (i.e., the equilibrium values got when the model equations were solved at MoI of 2 with the initial values of Lyt and Lys being equal to zero). That is, it was checked whether at MoI of 1 Lyt and Lys, starting from their equilibrium values at MoI of 2, could settle down to values which are representative of lysogeny (i.e., low values of Lyt and high values of Lys). It should be noted that this method finds out only those second stable states which represent lysogeny maintenance. Stochastic bistability at MoI of 1 was checked the same way. In order to ascertain the stability of lytic development, the model equations were solved at MoI of 2 with the initial values of Lyt and Lys being their equilibrium values at MoI of 1 (i.e., the equilibrium values got when the model equations were solved at MoI of 1 with the initial values of Lyt and Lys being equal to zero). That is, it was checked whether at MoI of 2 Lyt and Lys, starting from their equilibrium values at MoI of 1, could settle down to values which are representative of lysis (i.e., high values of Lyt and low values of Lys). As previously, the method finds out only those second stable states which are relevant for the stability of lytic development. Again stochastic bistability at MoI of 2 was checked the same way.

