## Supplementary Information for "Design Principles of Lambda’s Lysis*/*Lysogeny Decision vis-a-vis Multiplicity of Infection"

### Protein Equilibrium Level as a Function of the Gene Dosage in Negative Autoregulation

We show that in a negative autoregulatory loop, the protein equilibrium level cannot decrease as the gene copy number is increased.

$$\frac{dX}{dt} = \frac{mk_1}{1 + \frac{X^a}{K_{D1}}} - k_2 X \tag{1}$$

$$\bar{X} = \frac{mk_1/k_2}{1 + \frac{\bar{X}^a}{K_{D1}}} \tag{2}$$

Under the condition of very weak repression (i.e., $\bar{X}^a \ll K_{D1}$), the expression 2 becomes $\bar{X} = mk_1/k_2$. In this limit, equilibrium value of the protein rises proportionally to the gene dosage. Under the condition of very strong repression (i.e., $\bar{X}^a \gg K_{D1}$), the expression 2 becomes $\bar{X} = (mk_1 K_{D1}/k_2)^{(1/a+1)}$. In this limit, the dependence of equilibrium value of the protein on the gene dosage goes as $m^{(1/a+1)}$. This means that equilibrium value of the protein still increases with the gene dosage, albeit very slowly. When a is very large, $\bar{X}$ assumes the constant value of one irrespective of the gene dosage. The above analysis holds true for all values of the ratio between $\bar{X}$ and $K_{D1}$ because $\bar{X}(1 + \bar{X}^a/K_{D1})$ is a monotonous function.

**Table 1:** Average and standard deviation of deterministic and stochastic switch quotients for a given range of stochastic success rate for time length of 25.

| **Model** | **AVG Deterministic SQ (SD) (SSR[a] ⩾ 95)** | | **AVG Stochastic SQ (SD) (SSR ⩾ 95)** | |
|---|---|---|---|---|
| | a=2, b=2 | a=2, b=4 | a=2, b=2 | a=2, b=4 |
| 1A_Lyt_Lys | none | none | none | none |
| 1A_Lyt(1)_Lys | none | none | none | none |
| 1B_Lyt_Lys | none | none | none | none |
| 1B_Lyt(1)_Lys | none | 0.9304 (N/A) | none | 0.7662 (N/A) |
| 2_Lyt_Lys[b] | none | none | none | none |
| 3_Lyt_Lys | none | none | none | none |
| 4_Lyt_Lys | none | none | none | none |
| 6_Lyt_Lys | none | none | none | none |
| Lyt_Lys_CII | none | N/A | none | N/A |
| Lyt_Lys_CII(1) | none | N/A | none | N/A |
| Lyt(1)_Lys_CII(1) | none | N/A | none | N/A |
| Lyt_Lys_CII'[c] | none | N/A | none | N/A |

[a] SSR = Stochastic Success Rate

[b] after removing parameter sets which actually are that of 4_Lyt_Lys.

[c] Parameter sets selected for bistability at MoI of 2.

**Table 2:** Average and standard deviation of deterministic and stochastic switch quotients for a given range of stochastic success rate for time length of 25.

| **Model** | **AVG Deterministic SQ (SD) (95 >SSR[a] ⩾ 90)** | | **AVG Stochastic SQ (SD) (95 >SSR ⩾ 90)** | |
|---|---|---|---|---|
| | a=2, b=2 | a=2, b=4 | a=2, b=2 | a=2, b=4 |
| 1A_Lyt_Lys | none | none | none | none |
| 1A_Lyt(1)_Lys | none | none | none | none |
| 1B_Lyt_Lys | none | 0.9270 (N/A) | none | 0.7855 (N/A) |
| 1B_Lyt(1)_Lys | none | none | none | none |
| 2_Lyt_Lys | none | none | none | none |
| 3_Lyt_Lys | none | none | none | none |
| 4_Lyt_Lys | none | none | none | none |
| 6_Lyt_Lys | none | none | none | none |
| Lyt_Lys_CII | 0.9816 (N/A) | N/A | 0.7916 (N/A) | N/A |
| Lyt_Lys_CII(1) | 0.9593 (N/A) | N/A | 0.6789 (N/A) | N/A |
| Lyt(1)_Lys_CII(1) | none | N/A | none | N/A |
| Lyt_Lys_CII'[c] | none | N/A | none | N/A |

[a] SSR = Stochastic Success Rate

**Table 3:** Average and standard deviation of deterministic and stochastic switch quotients for a given range of stochastic success rate for time length of 50.

| **Model** | **AVG Deterministic SQ (SD) (SSR[a] ⩾ 95)** | | **AVG Stochastic SQ (SD) (SSR ⩾ 95)** | |
|---|---|---|---|---|
| | a=2, b=2 | a=2, b=4 | a=2, b=2 | a=2, b=4 |
| 1A_Lyt_Lys | none | none | none | none |
| 1A_Lyt(1)_Lys | none | none | none | none |
| 1B_Lyt_Lys | none | none | none | none |
| 1B_Lyt(1)_Lys | none | 0.9304 (N/A) | none | 0.7362 (N/A) |
| 2_Lyt_Lys[b] | none | none | none | none |
| 3_Lyt_Lys | none | none | none | none |
| 4_Lyt_Lys | none | none | none | none |
| 6_Lyt_Lys | none | none | none | none |
| Lyt_Lys_CII | none | N/A | none | N/A |
| Lyt_Lys_CII(1) | 0.9718 (N/A) | N/A | 0.7411 (N/A) | N/A |
| Lyt(1)_Lys_CII(1) | none | N/A | none | N/A |
| Lyt_Lys_CII'[c] | none | N/A | none | N/A |

[a] SSR = Stochastic Success Rate
[b] after removing parameter sets which actually are that of 4_Lyt_Lys.
[c] Parameter sets selected for bistability at MoI of 2.

**Table 4:** Average and standard deviation of deterministic and stochastic switch quotients for a given range of stochastic success rate for time length of 50.

| **Model** | **AVG Deterministic SQ (SD) (95 >SSR[a] $\geqslant$ 90)** | | **AVG Stochastic SQ (SD) (95 >SSR $\geqslant$ 90)** | |
|---|---|---|---|---|
| | a=2, b=2 | a=2, b=4 | a=2, b=2 | a=2, b=4 |
| 1A_Lyt_Lys | 0.9928 (0.0031) | 0.9946 (0.0009) | 0.8392 (0.0181) | 0.6894 (0.0291) |
| 1A_Lyt(1)_Lys | none | none | none | none |
| 1B_Lyt_Lys | 0.9961 (0.0010) | 0.9270 (N/A) | 0.8226 (0.0135) | 0.7924 (N/A) |
| 1B_Lyt(1)_Lys | none | none | none | none |
| 2_Lyt_Lys | none | none | none | none |
| 3_Lyt_Lys | none | none | none | none |
| 4_Lyt_Lys | none | 0.4956 (0.0006) | none | 0.3194 (0.0044) |
| 6_Lyt_Lys | none | none | none | none |
| Lyt_Lys_CII | 0.9729 (0.0110) | N/A | 0.7530 (0.0190) | N/A |
| Lyt_Lys_CII(1) | 0.9671 (0.0124) | N/A | 0.7866 (0.0343) | N/A |
| Lyt(1)_Lys_CII(1) | none | N/A | none | N/A |
| Lyt_Lys_CII'[c] | none | N/A | none | N/A |

[a] SSR = Stochastic Success Rate

**Table 5:** Average and standard deviation of deterministic and stochastic switch quotients for a given range of stochastic success rate for time length of 200.

| **Model** | **AVG Deterministic SQ (SD) (SSR[a] $\geqslant$ 95)** | | **AVG Stochastic SQ (SD) (SSR $\geqslant$ 95)** | |
|---|---|---|---|---|
| | a=2, b=2 | a=2, b=4 | a=2, b=2 | a=2, b=4 |
| 1A_Lyt_Lys | 0.9923 (0.0023) | 0.9896 (0.0056) | 0.7591 (0.0568) | 0.5830 (0.0877) |
| 1A_Lyt(1)_Lys | none | 0.9952 (0.0022) | none | 0.5548 (0.0211) |
| 1B_Lyt_Lys | 0.9961 (0.0010) | 0.9855 (0.0262) | 0.7908 (0.0294) | 0.6694 (0.0861) |
| 1B_Lyt(1)_Lys | 0.9469 (N/A) | 0.9643 (0.0339) | 0.4660 (N/A) | 0.7275 (0.0420) |
| 2_Lyt_Lys[b] | none | none | none | none |
| 3_Lyt_Lys | none | none | none | none |
| 4_Lyt_Lys | none | 0.4956 (0.0006) | none | 0.2904 (0.0117) |
| 6_Lyt_Lys | none | none | none | none |
| Lyt_Lys_CII | 0.9729 (0.0110) | N/A | 0.7677 (0.03441) | N/A |
| Lyt_Lys_CII(1) | 0.9655 (0.0062) | N/A | 0.7406 (0.0139) | N/A |
| Lyt(1)_Lys_CII(1) | 0.9647 (N/A) | N/A | 0.7150 (N/A) | N/A |
| Lyt_Lys_CII'[c] | 0.9772 (0.0098) | N/A | 0.6076 (0.0198) | N/A |

[a] SSR = Stochastic Success Rate
[b] after removing parameter sets which actually are that of 4_Lyt_Lys.
[c] Parameter sets selected for bistability at MoI of 2.

**Table 6:** Average and standard deviation of deterministic and stochastic switch quotients for a given range of stochastic success rate for time length of 200.

| **Model** | **AVG Deterministic SQ (SD) (95 >SSR[a] ⩾ 90)** | | **AVG Stochastic SQ (SD) (95 >SSR ⩾ 90)** | |
|---|---|---|---|---|
| | a=2, b=2 | a=2, b=4 | a=2, b=2 | a=2, b=4 |
| 1A_Lyt_Lys | 0.9853 (0.0191) | 0.9886 (0.0043) | 0.7850 (0.0265) | 0.3989 (0.0511) |
| 1A_Lyt(1)_Lys | 0.9945 (N/A) | 0.9904 (0.0048) | 0.8031 (N/A) | 0.4932 (0.0526) |
| 1B_Lyt_Lys | 0.9727 (0.0256) | 0.9679 (0.0239) | 0.7782 (0.0283) | 0.4621 (0.0125) |
| 1B_Lyt(1)_Lys | 0.9826 (0.0270) | 0.9898 (0.0078) | 0.7148 (0.0504) | 0.5115 (0.0585) |
| 2_Lyt_Lys | none | none | none | none |
| 3_Lyt_Lys | none | 0.9802 (0.0206) | none | 0.5482 (0.0423) |
| 4_Lyt_Lys | none | none | none | none |
| 6_Lyt_Lys | none | none | none | none |
| Lyt_Lys_CII | none | N/A | none | N/A |
| Lyt_Lys_CII(1) | 0.9936 (N/A) | N/A | 0.8413 (N/A) | N/A |
| Lyt(1)_Lys_CII(1) | none | N/A | none | N/A |
| Lyt_Lys_CII'[c] | 0.9786 (0.0116) | N/A | 0.6260 (0.0709) | N/A |

[a] SSR = Stochastic Success Rate

**Table 7:** Number of parameter sets for various ranges of stochastic success rate for time length 25.

| Model | SSR[a] ⩾ 95 | | 95 >SSR ⩾ 90 | | 90 >SSR ⩾ 80 | | Total no. of parameter sets | |
|---|---|---|---|---|---|---|---|---|
| | a=2 b=2 | a=2 b=4 | a=2 b=2 | a=2 b=4 | a=2 b=2 | a=2 b=4 | a=2 b=2 | a=2 b=4 |
| 1A_Lyt_Lys | 0 | 0 | 0 | 0 | 8 | 3 | 20 | 17 |
| 1A_Lyt(1)_Lys | 0 | 0 | 0 | 0 | 1 | 0 | 17 | 15 |
| 1B_Lyt_Lys | 0 | 0 | 0 | 1 | 2 | 2 | 6 | 11 |
| 1B_Lyt(1)_Lys | 0 | 1 | 0 | 0 | 1 | 0 | 14 | 10 |
| 2_Lyt_Lys | 0 | 0 | 0 | 0 | 0 | 0 | 1 | 6 |
| 3_Lyt_Lys | 0 | 0 | 0 | 0 | 0 | 0 | 10 | 12 |
| 4_Lyt_Lys | 0 | 0 | 0 | 0 | 0 | 2 | 2 | 2 |
| 6_Lyt_Lys | 0 | 0 | 0 | 0 | 0 | 0 | 1 | 13 |
| Lyt_Lys_CII | 0 | N/A | 1 | N/A | 3 | N/A | 9 | N/A |
| Lyt_Lys_CII(1) | 0 | N/A | 1 | N/A | 4 | N/A | 9 | N/A |
| Lyt(1)_Lys_CII(1) | 0 | N/A | 0 | N/A | 2 | N/A | 9 | N/A |
| Lyt_Lys_CII'[c] | 0 | N/A | 0 | N/A | 2 | N/A | 9 | N/A |

[a] SSR = Stochastic Success Rate

**Table 8:** Number of parameter sets for various ranges of stochastic success rate for time length 50.

| Model | SSR[a] ⩾ 95 | | 95 >SSR ⩾ 90 | | 90 >SSR ⩾ 80 | | Total no. of parameter sets | |
|---|---|---|---|---|---|---|---|---|
| | a=2 b=2 | a=2 b=4 | a=2 b=2 | a=2 b=4 | a=2 b=2 | a=2 b=4 | a=2 b=2 | a=2 b=4 |
| 1A_Lyt_Lys | 0 | 0 | 3 | 2 | 10 | 5 | 20 | 17 |
| 1A_Lyt(1)_Lys | 0 | 0 | 0 | 0 | 3 | 2 | 17 | 15 |
| 1B_Lyt_Lys | 0 | 0 | 2 | 1 | 1 | 4 | 6 | 11 |
| 1B_Lyt(1)_Lys | 0 | 1 | 0 | 0 | 2 | 1 | 14 | 10 |
| 2_Lyt_Lys | 0 | 0 | 0 | 0 | 0 | 0 | 1 | 6 |
| 3_Lyt_Lys | 0 | 0 | 0 | 0 | 1 | 0 | 10 | 12 |
| 4_Lyt_Lys | 0 | 0 | 0 | 2 | 0 | 0 | 2 | 2 |
| 6_Lyt_Lys | 0 | 0 | 0 | 0 | 0 | 0 | 1 | 13 |
| Lyt_Lys_CII | 0 | N/A | 3 | N/A | 1 | N/A | 9 | N/A |
| Lyt_Lys_CII(1) | 1 | N/A | 3 | N/A | 2 | N/A | 9 | N/A |
| Lyt(1)_Lys_CII(1) | 0 | N/A | 0 | N/A | 2 | N/A | 9 | N/A |
| Lyt_Lys_CII'[c] | 0 | N/A | 0 | N/A | 3 | N/A | 9 | N/A |

[a] SSR = Stochastic Success Rate

**Table 9:** Number of parameter sets for various ranges of stochastic success rate for time length 200.

| Model | SSR[a] ⩾ 95 | | 95 >SSR ⩾ 90 | | 90 >SSR ⩾ 80 | | Total no. of parameter sets | |
|---|---|---|---|---|---|---|---|---|
| | a=2 b=2 | a=2 b=4 | a=2 b=2 | a=2 b=4 | a=2 b=2 | a=2 b=4 | a=2 b=2 | a=2 b=4 |
| 1A_Lyt_Lys | 8 | 7 | 5 | 5 | 4 | 4 | 20 | 17 |
| 1A_Lyt(1)_Lys | 0 | 2 | 1 | 6 | 5 | 6 | 17 | 15 |
| 1B_Lyt_Lys | 2 | 6 | 2 | 2 | 0 | 2 | 6 | 11 |
| 1B_Lyt(1)_Lys | 1 | 2 | 4 | 5 | 4 | 2 | 14 | 10 |
| 2_Lyt_Lys | 0 | 0 | 0 | 0 | 0 | 0 | 1 | 6 |
| 3_Lyt_Lys | 0 | 0 | 0 | 4 | 1 | 6 | 10 | 12 |
| 4_Lyt_Lys | 0 | 2 | 0 | 0 | 0 | 0 | 2 | 2 |
| 6_Lyt_Lys | 0 | 0 | 0 | 0 | 0 | 2 | 1 | 13 |
| Lyt_Lys_CII | 3 | N/A | 0 | N/A | 0 | N/A | 9 | N/A |
| Lyt_Lys_CII(1) | 2 | N/A | 1 | N/A | 4 | N/A | 9 | N/A |
| Lyt(1)_Lys_CII(1) | 1 | N/A | 0 | N/A | 1 | N/A | 9 | N/A |
| Lyt_Lys_CII′[c] | 3 | N/A | 3 | N/A | 0 | N/A | 9 | N/A |

[a] SSR = Stochastic Success Rate

**Table 10:** The parameter set selected for 2_Lyt_Lys but which actually defines 4_Lyt_Lys for a=2, b=4.

| Model | avg $k_1$ (SD) | avg $k_3$ (SD) | avg $K_{D1}$ (SD) | avg $K_{D2}$ (SD) |
|---|---|---|---|---|
| 2_Lyt_Lys[a] | 15.01 (N/A) | 5.00 (N/A) | 38032.88 (N/A) | 1240.42 (N/A) |
| 2_Lyt_Lys[b] | 10.00 (0) | 5.01 (0.21) | 0.35 (0.30) | 0.02 (0.02) |
| 4_Lyt_Lys | 20.08 (2.50) | 5.02 (≈ 0) | N/A | 672.23 (167.92) |

[a] parameter set which actually is that of 4_Lyt_Lys
[b] Rest of the parameter sets

**Table 11:** The parameter set selected for 6_Lyt_Lys but which actually defines 2_Lyt_Lys for a=2, b=4.

| **Model** | **avg $k_1$ (SD)** | **avg $k_3$ (SD)** | **avg $k_4$ (SD)** | **avg $K_{D1}$ (SD)** | **avg $K_{D2}$ (SD)** |
|---|---|---|---|---|---|
| 6_Lyt_Lys[a] | 10.00 (0) | 11.28 (7.90) | 4.94 (0.14) | 0.1180 (0.0654) | 0.0159 (0.0105) |
| 6_Lyt_Lys[b] | 10.00 (0) | 199.41 (90.16) | 4.59 (0.13) | 0.0764 (0.0437) | 21.85 (4.02) |
| 6_Lyt_Lys[c] | 10.00 (0.0056) | 0.23 (0.33) | 4.95 (0.09) | 137.46 (312.32) | 0.004 (0.007) |

[a] parameter sets which actually define 2_Lyt_Lys but with $k_3$ of two of the sets being higher (20.08 and 8.75 vs 5.02)
[b] (three) parameter sets whose stochastic success rate is either zero or negligible (see main text)
[c] Rest of the parameter sets

**Table 12:** Parameter sets of Lyt_Lys_CII with and without bistability at MoI of 2.

| **Bistability at MoI of 2** | **avg $k_1$ (SD)** | **avg $k_2$ (SD)** | **avg $k_3$ (SD)** | **avg $k_4$ (SD)** | **avg $k_5$ (SD)** | **avg $K_{D1}$ (SD)** | **avg $K_{D2}$ (SD)** | **avg $K_{D3}$ (SD)** |
|---|---|---|---|---|---|---|---|---|
| No | 18.40 (6.00) | 0.5862 (0.1738) | 0.1537 (0.0562) | 0.1990 (0.1832) | 5.1040 (0.0867) | 2541.55 (646.82) | 0.6525 (0.1978) | 20.98 (23.97) |
| Yes | 18.99 (7.58) | 0.5823 (0.2959) | 0.1109 (0.1205) | 0.0814 (0.1663) | 5.003 (0.0804) | 192.04 (202.73) | 0.0439 (0.0447) | 0.0629 (0.1431) |
